# The geometry of the representation of decision variable and stimulus difficulty in the parietal cortex

**DOI:** 10.1101/2021.01.04.425244

**Authors:** Gouki Okazawa, Christina E. Hatch, Allan Mancoo, Christian K. Machens, Roozbeh Kiani

**Affiliations:** Center for Neural Science, New York University, New York, NY 10003, USA; Champalimaud Research, Champalimaud Centre for the Unknown, 1400-038 Lisbon, Portugal; Neuroscience Institute, NYU Langone Medical Center, New York, NY 10016, USA; Department of Psychology, New York University, New York, NY 10003, USA

## Abstract

Lateral intraparietal (LIP) neurons represent formation of perceptual decisions involving eye movements. In circuit models for these decisions, neural ensembles that encode actions compete to form decisions. Consequently, decision variables (DVs) are represented as partially potentiated action plans, where ensembles increase their average responses for stronger evidence supporting their preferred actions. As another consequence, DV representation and readout are implemented similarly for decisions with identical competing actions, irrespective of input and task context differences. Here, we challenge those core principles using a novel face-discrimination task, where LIP firing rates decrease with supporting evidence, contrary to conventional motion-discrimination tasks. These opposite response patterns arise from similar mechanisms in which decisions form along curved population-response manifolds misaligned with action representations. These manifolds rotate in state space based on task context, necessitating distinct readouts. We show similar manifolds in lateral and medial prefrontal cortices, suggesting a ubiquitous representational geometry across decision-making circuits.

## Introduction

Perceptual decision making relies on deliberative processes that evaluate and compare possible alternatives based on sensory information. A commonly suggested mechanism for such decisions is to integrate sensory information into a decision variable (DV) and compare it against a decision criterion or bound to commit to a choice (Green and Swets 1966; Link 1992; Ratcliff and McKoon 2008; Smith and Vickers 1988). Consistent with these behavioral models, multiple brain structures show activity compatible with representation of the DV (Gold and Shadlen 2007; Shadlen and Kiani 2013). For example, neurons in the lateral intraparietal (LIP) area increase their average firing rate in response to sensory evidence supporting their preferred saccade target during a motion direction discrimination task (Shadlen and Newsome 2001). The idea that average LIP firing rate reflects accumulated evidence has successfully accounted for various aspects of decision-making behavior including choice and reaction time distributions (Churchland et al. 2008; Purcell and Kiani 2016), choice confidence (Kiani and Shadlen 2009), and adjustments of decision criteria with prior probabilities (Hanks et al. 2011; Rao et al. 2012), reward imbalance (Rorie et al. 2010), speed-accuracy instructions (Hanks et al. 2014), and feedback-dependent changes of future decisions (Purcell and Kiani 2016). Similar encoding of the DV has been found in many other brain structures (de Lafuente et al. 2015; Deverett et al. 2018; Ding and Gold 2010, 2012; Donner et al. 2009; Horwitz and Newsome 1999; Kiani et al. 2014b; Kim and Basso 2008; Kim and Shadlen 1999; Peixoto et al. 2018; Ratcliff et al. 2003) and in other perceptual tasks (Hanks et al. 2015; Heekeren et al. 2004; O’Connell et al. 2012; Philiastides and Sajda 2006; Purcell et al. 2010; Thura and Cisek 2014).

These experimental findings have promoted the proposition that decisions are formed through competition of neuronal modules with specific choice preferences until one module “wins” and dictates the choice. The strongest supporting evidence for this proposition is observed in motor planning regions of the brain, where neurons represent the decision-making process when a decision is communicated through their encoding actions. Sensory information biases this competition such that the evidence supporting a choice increases the activity of corresponding neurons. Since the competition happens at the level of action selection, what distinguishes different perceptual tasks is the sensory information being integrated, not the integrator itself. These ideas shape existing theoretical frameworks for implementation of the decision-making process with biophysically-realistic neural networks (Beck et al. 2008; Deco et al. 2013; Lo and Wang 2006; Mazurek et al. 2003; Purcell et al. 2010; Wang 2002; Wimmer et al. 2015; Wong and Wang 2006). Such networks have been quite successful in explaining past behavioral and physiological data.

However, several key assumptions at the core of these frameworks have yet to be tested experimentally. First, it is unclear whether response patterns that encode the DVs are explainable as intermediate patterns between competing actions, and whether the pools of neurons that represent the competing actions are also the ones that compete during decision formation. These neurons clearly represent the decision-making process when a decision is associated with their encoding action, but the contentious point is whether this representation is inseparably tied to the choice or action selectivity of these neurons. Second, it is unclear whether the neural representation of the integration process is shared across tasks and whether potential differences necessitate different readout schemes for downstream circuits. Testing these assumptions has been challenging as past studies were largely limited to a single behavioral task and focused on average responses across the recorded neurons. A side by side comparison of multiple tasks is necessary for a rigorous test as it creates an opportunity to observe qualitative and quantitative differences of neural activity across tasks. Further, proper comparisons across tasks necessitate looking beyond simple averages and exploring population response patterns, as those patterns could include properties obscured by averaging and not immediately obvious from single neuron responses (Cunningham and Yu 2014; Raposo et al. 2014; Rigotti et al. 2013).

Here we examine the geometry of population activity in LIP in seven monkeys and three tasks: a motion direction discrimination task, which has been extensively used to study perceptual decisions, and two variants of a novel face categorization task, where subjects report the species or expression of a face (Okazawa et al. 2018). We find a striking mismatch in patterns of average firing rates among the tasks. Specifically, mean firing rates during the face tasks are inversely correlated with the sensory evidence in favor of their preferred saccade targets — the opposite of the pattern observed in the motion task (Churchland et al. 2008; Kiani et al. 2008; Shadlen and Newsome 2001). We show that this inconsistency arises because, at each moment, the LIP population encodes the DV on a curved manifold in the state space and this manifold changes in a task-dependent manner. We establish that the curved manifold provides an explicit code (Misaki et al. 2010) for both the DV and stimulus difficulty, but the latter does not have apparent functional contributions to the monkey’s confidence about the choice. Further, we demonstrate that task-dependent curvature of the manifold significantly undermines the hypothesis that decisions are formed through competition of action-selective modules. Rather, decisions are initially formed in a space dissociated from action encoding and then translated to an action plan at later stages of decision making. By exploring species and expression categorization of the same faces while recording from the same neurons, we show that task dependence of curved manifolds is not caused solely by differences of sensory stimuli; they are also strongly shaped by task requirements. Finally, we demonstrate that similar curved manifolds emerge in the lateral and medial prefrontal cortices and are not limited to the LIP. We propose that the curvature of integration manifold is a fundamental representational property of the neural circuits encoding the DV. Our findings invite a major update in existing computational frameworks of perceptual decision making.

## Results

### Curved population response manifolds in LIP

We examined population response properties in LIP for two variants of a face categorization task (Fig. 1A, face task) and a motion direction discrimination task (Fig. 1B, motion task). In the motion task, monkeys viewed a dynamic random-dots stimulus and reported the perceived motion direction. We manipulated stimulus difficulty by varying the percentage of coherently moving dots across trials (motion coherence). Within each trial, the stochastic nature of the stimulus caused frame by frame fluctuations of motion coherence around a nominal mean for the trial. In the face task, monkeys classified the species of a set of face stimuli (monkey or human) or their expression (sad or happy) in separate blocks. The discriminated categories in each block were defined by two prototype faces. We manipulated stimulus difficulty by creating a morph continuum between the two prototypes and varying the stimulus morph level across trials. Our customized morphing algorithm allowed independent morphing of facial features. We chose eyes, nose, and mouth as informative features and morphed them, while fixing other facial features at halfway between the prototypes. The facial features fluctuated randomly around a mean morph level in each trial (SD, 20% morph; updated every 106.7 ms), similar to the variability of motion coherence during each trial of the motion task. These fluctuations, which were interleaved with masks to remain subliminal, enabled us to estimate spatiotemporal weighting of informative features for decision making (Fig. S1). In both tasks, monkeys reported their decision by making a saccadic eye movement to one of the two targets, one placed inside the response field (RF) of the recorded neurons (T_in_) and the other on the opposite side of the screen, outside the RF (T_out_).

**Figure 1:**
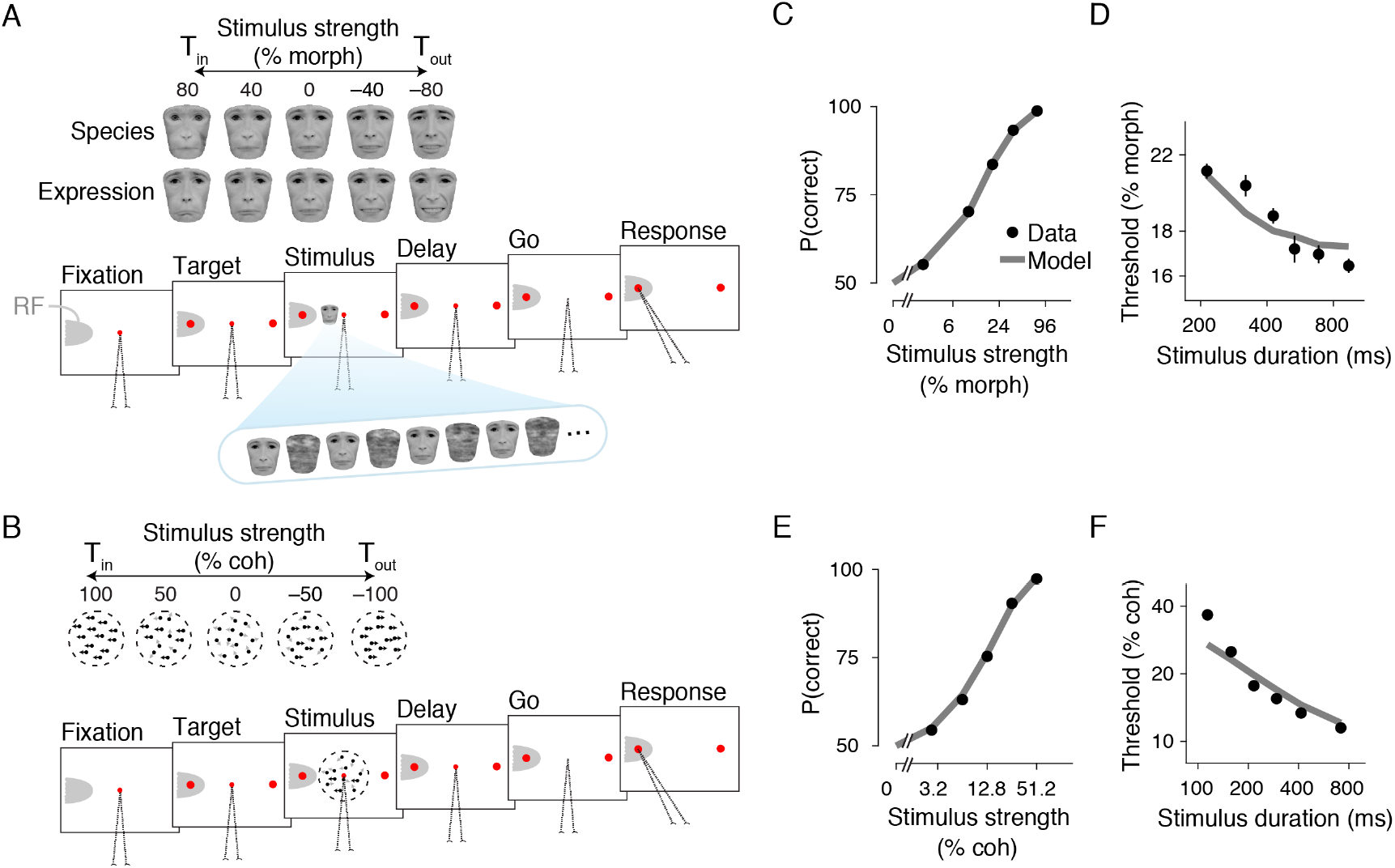
Task designs and behavioral results. (**A**) Face categorization task. Monkeys classified face stimuli, reporting either the species (monkey vs. human) or expression (sad vs. happy) of the faces in different blocks. Stimuli in each session were chosen from a morph plane defined by a monkey and a human face (prototypes for species categorization), and sad and happy expressions (prototypes for expression categorization). The two morph axes from an example session are shown at the top of the panel. On each trial, after the monkey gazed at a central fixation point (FP), two target dots appeared, one inside (T_in_) and the other outside (T_out_) of the neuron RF. Then, a sequence of faces interleaved with masks appeared on the screen. During the sequence, three facial features (eyes, nose, mouth) fluctuated around a nominal mean morph level for the trial. Other facial features remained constant across the block and were uninformative. The changing features in a trial enabled us to estimate the weights of different features for the decision-making process (Fig. S1). The masks prevented change detection of facial features within the trial, making the stimulus appear as a naturalistic face covered with varying cloudy patterns (Okazawa et al. 2020). The stimulus was positioned non-foveally and its size was randomly varied across trials by one octave to prevent the monkey from relying on local and low-level visual attributes. Stimulus presentation was followed by a delay period. At the end of the delay, the FP disappeared (Go cue), and the monkey reported its choice by making a saccade to one of the targets. The displayed human face images were obtained from Radboud database (Langner et al. 2010) and used with permission for visualization purpose. (**B**) Motion direction discrimination task. Monkeys viewed a dynamic random-dots motion and reported the net motion direction. Stimulus difficulty was controlled by varying the percentage of coherently moving dots on each trial. The sequence of task events was similar to that of the face task. (**C, E**) In both the face (C) and the motion (E) tasks, monkeys’ accuracy monotonically improved for higher stimulus strengths. Gray lines are the fits of drift diffusion models (see Methods). (**D, F**) Psychophysical thresholds decreased for longer stimulus durations. The stimulus strength corresponding to 81.6% accuracy was defined as a psychophysical threshold. Error bars are s.e.m.

Three monkeys performed the motion task and two monkeys performed the face task, while we recorded from LIP neurons. In both tasks, the monkey’s choice accuracy monotonically improved as a function of stimulus strength (Fig. 1C, E) and duration (Fig. 1D, F). The reduction of psychophysical thresholds with longer stimulus durations (face task, log-log regression slope, −0.19 ± 0.02, *p* < 0.001; motion task, −0.61 ± 0.03, *p* < 0.001, bootstrap test) indicated that monkeys leveraged multiple stimulus frames to make their decisions. These results were quantitatively compatible with drift diffusion models (DDMs) that accumulated sensory evidence toward decision bounds in both tasks (DDM fit to psychometric function, face task, *R*^2^ = 0.998; motion task, *R*^2^ = 0.997). In both tasks the model also quantitatively explained the results of psychophysical reverse correlation, including differential effects of sensory evidence on choice over time, and differential weighting of informative facial features for species and expression categorizations (Fig. S1). Based on the similarity of decision-making behavior between the two tasks, we hypothesized that LIP neurons would exhibit similar decision-related responses in both tasks.

Contrary to our expectation, we found a striking difference in the average LIP responses in the two tasks. Fig. 2A-B shows the population average peri-stimulus time histograms (PSTHs) after stimulus onset, plotted separately for choosing the T_in_ (solid lines) and T_out_ (dashed lines). In the motion task (Fig. 2B), neurons had higher firing rates when the stimulus strongly supported the T_in_ choice and lower firing rates when it strongly supported the T_out_ choice. Thus, their firing rates monotonically reflected the amount of evidence for the T_in_ choice. This pattern has been consistently reported in the past for the motion task (Kumano et al. 2016; Rao et al. 2012; Roitman and Shadlen 2002; Rorie et al. 2010; Shadlen and Newsome 2001; Shushruth et al. 2018) and prompted the conclusion that the average firing rates represent the DV and the propensity to make a particular choice. However, LIP neurons recorded in the face task exhibited a reversed order for T_in_ choices (Fig. 2A); they showed enhanced activity for weaker sensory evidence. Consequently, the average firing rates no longer monotonically reflected the amount of evidence for the T_in_ choice. This observation is puzzling as it seemingly contradicts the previous account of LIP responses as a neural correlate of accumulated evidence for a decision (Mazurek et al. 2003). Critically, this pattern also appears to disagree with monkeys’ behavioral performance, which monotonically improved as a function of stimulus strength in both tasks (Fig. 1).

**Figure 2:**
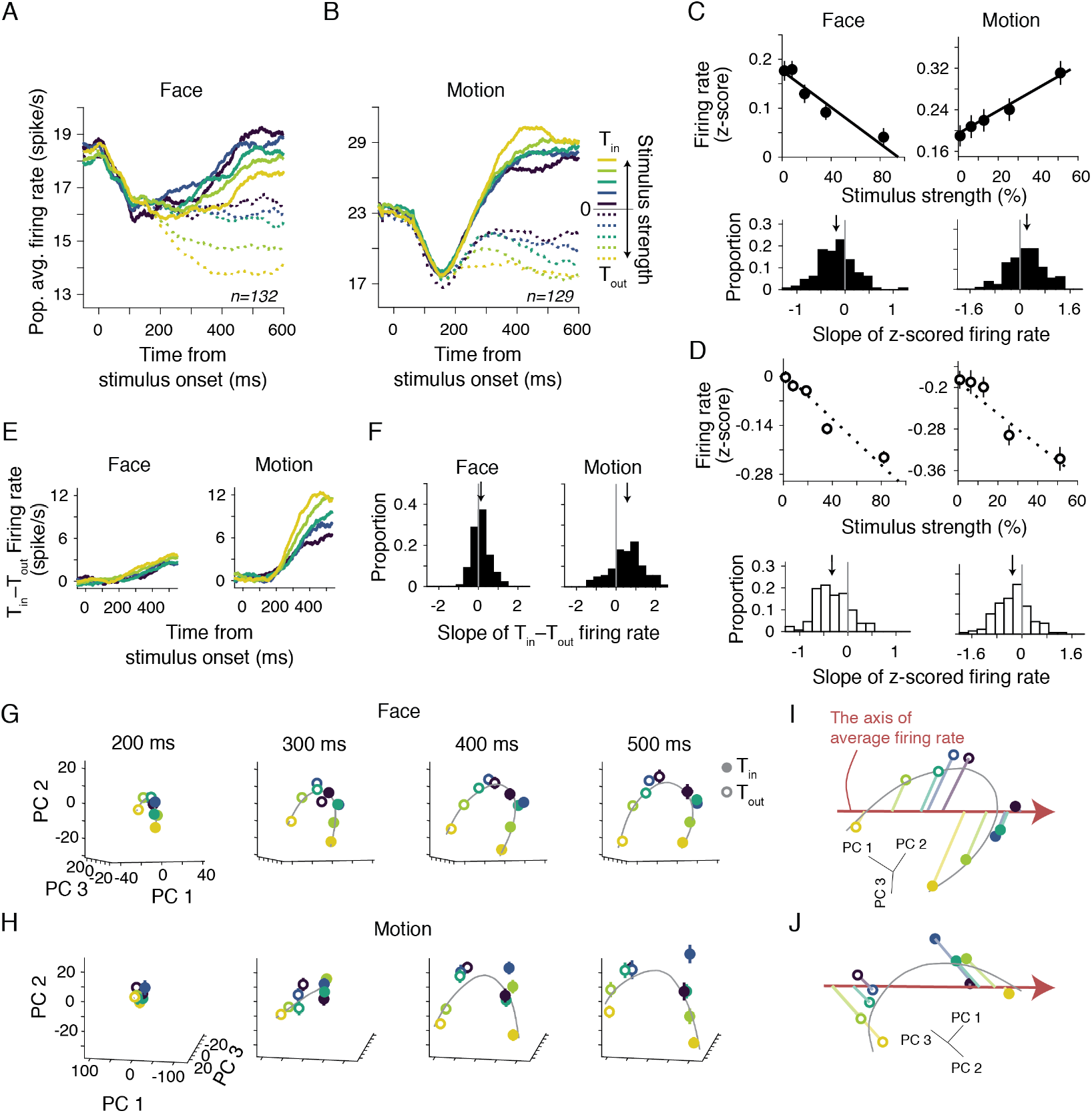
Population response patterns form a curved manifold in state space that rotates across tasks, giving rise to the opposite ordering of population average PSTHs in the two tasks. (**A, B**) Population-averaged PSTHs showed opposite ordering of firing rates as a function of stimulus strength for T_in_ choices (solid lines) in the face (A) and motion (B) tasks. Stronger motion toward T_in_ was associated with higher firing rates, matching past studies (Kumano et al. 2016; Roitman and Shadlen 2002; Shadlen and Newsome 2001; Shushruth et al. 2018). The face task had a reversed order. The PSTHs for T_out_ choices (dashed lines) showed a similar order in the two tasks. Only correct choices were included here to ensure that the order of PSTHs was not shaped by different proportions of choices for different stimulus strengths. But the same order was also present in the PSTHs unconditioned on choice (Fig. S2E). (**C**) Firing rates (250–600 ms window) as a function of stimulus strength for T_in_ choices. The average firing rates increased with the stimulus strength in the motion task but decreased in the face task (top). Histograms of the slope of individual unit firing rates as a function of stimulus strength (*α*_1_ in Eq.1) illustrate the prevalence of the two trends (bottom). Arrows indicate means. (**D**) Firing rates and slope histograms for T_out_ choices. (**E**) The difference of T_in_ and T_out_ firing rates were larger for stronger stimuli in the motion task but almost identical for different stimulus strengths in the face task. (**F**) Slope histograms for the difference of T_in_ and T_out_ firing rates as a function of stimulus strength (*β*_1_ in Eq.2). (**G**) Population neural responses formed a curved manifold for the face task. Principal component analysis (PCA) was performed on trial-averaged PSTHs concatenated across units. The panel depicts neural responses projected onto a 3D state space formed by the top three PCs. The PC axes were derived using PSTHs of 250-600 ms window. All panels show the same PC space. The points correspond to the population neural activity for correct choices to different stimulus strengths at different times after stimulus onset. The gray lines are cubic smoothing splines fit to the data. (**H**) Same as G but for the motion task. (**I, J**) The relationship between average firing rates and population response manifolds. Changes in the population average firing rate correspond to a linear axis in the PC space (red vector). Projection of the manifold onto this axis determines the population average firing rates for different stimulus strengths. Depending on the direction of manifold curvature, these projections could lead to monotonic (motion task, J) or non-monotonic (face task, I) changes of population average activity as a function of stimulus strength. The projection lines are perpendicular to the axis of average rates in the state space, though they appear non-perpendicular in 2D illustrations. Error bars are s.e.m.

To quantify the significance of this reversal in firing rates, we plotted the population average firing rates in a 250-600 ms window from stimulus onset as a function of stimulus strength (Fig. 2C). For T_in_ choices in the motion task, firing rates increased with the strength of evidence supporting the choice (*α*_1_ = 0.22 ± 0.05 across units, *p* = 8.2 × 10^−5^,; Eq.1), whereas in the face task, firing rates decreased (*α*_1_ = −0.19 ± 0.03, *p* = 3.8 × 10^−8^). This reversal was clearly seen in many single units and was consistently present in different monkeys and different task variations (Fig. S2A-C). Later in the trial, especially immediately before T_in_ saccades, these differential responses disappeared and the firing rates reached a common level regardless of stimulus strength (Fig. S4A), as has been reported previously (Shadlen and Newsome 2001). In contrast to T_in_ choices, firing rates systematically declined with the strength of evidence in favor of the T_out_ choice during stimulus viewing in both tasks (Fig. 2D; motion task, *α*_1_ = −0.30 ± 0.05, *p* = 5.0 × 10^−9^; face task, *α*_1_ = −0.33 ± 0.03, *p* = 2.9 × 10^−20^). The PSTHs for the T_out_ choice were better separated for different stimulus strengths (Fig. 2B), consistent with past electrophysiological studies (Shadlen and Newsome 2001) as well as bounded evidence accumulation models for the decision-making process.

If a decision is formed through competition between neurons that encode the two choices, then the difference of firing rates associated with the T_in_ and T_out_ choices would be larger for the stronger stimuli. This was indeed the case in the motion task (Fig. 2E-F; the slope of the differential firing rates as a function of stimulus strength: *β*_1_ = 0.57 ± 0.07; Eq.2). However, the T_in_−T_out_ differential firing rates were much less dependent on the stimulus strength in the face task (*β*_1_ = 0.13 ± 0.04; difference from the motion task: *p* = 8.1 × 10^−8^) due to the reversal of T_in_ firing rates. Because of these response properties, we can expect that the difference of activity of pools of neurons selective for the rightward and leftward saccades would fail to explain the higher accuracy of behavior for stronger face stimuli. To illustrate this point, we used a receiver operating characteristic analysis to quantify how well an ideal observer could discriminate the stimuli supporting the two choices if it could monitor responses of a LIP neuron (and its anti-neuron partner) (Britten et al. 1992). The ideal observer had systematically higher accuracy for stronger stimuli in the motion task (Figure S2D), especially after 300ms from stimulus onset where T_in_ and T_out_ PSTHs diverge well. But the same ideal observer did not benefit from stronger stimuli in the face task because the difference of T_in_ and T_out_ PSTHs remains largely independent of stimulus strength.

Does the reversal suggest different underlying neural computations during face categorization? Because individual units show diverse response profiles (see the distributions in Fig. 2C-D), we sought to better understand the neural code by exploring the patterns of population responses in the two tasks. Figure 2G-H depicts population activity in a three-dimensional (3D) principal component (PC) space derived from the PSTHs of recorded units (top 3 PCs, explained variance, motion task: 78%, face task: 54%). The data points in this state space correspond to the population responses for different stimulus strengths at different times after stimulus onset. The population neural responses gradually changed along a curved line in the state space that connected the strongest stimulus supporting T_in_ to the strongest stimulus supporting T_out_ choices. In both tasks, the curvature was formed such that the distance from the base of the curve correlated with the unsigned stimulus strength. This curvature was quite large and unlikely to have been caused by random fluctuation of neural responses. Specifically, a bootstrap analysis confirmed that the manifold is consistently curved along one direction (*p* < 0.001 for both tasks, see Methods for details; see also Fig. S5A for an alternative test of the significance of curvature). This curvature was observed in individual monkeys and also found for groups of simultaneously recorded units within single sessions (Fig. S5B-D), confirming that it is a robust feature of LIP population responses during decision formation. This curvature arises from the diverse tuning of neurons for stimulus strength across the population (Fig. S3B). The curved manifold that spans different stimulus strengths gradually expanded over time after stimulus onset, indicating better separation of population response patterns associated with different stimuli. This expansion is compatible with integration of sensory evidence over time and the representation of the DV by the neural population. However, later in the trial and especially close to the saccade initiation, the population response patterns converged to two distinct points in state space that reflected the monkey’s choices (Fig. S4B-D), suggesting replacement of the representation of the DV with the representation of choice.

An important implication of the curvature of the population response manifold is that the *average* firing rate does not necessarily preserve the order of the stimulus strengths. Recall that the average firing rate is calculated as a linear sum of single-neuron firing rates with equal weights across the recorded population. Since projections on the PC space are also calculated as a linear weighted sum of single-neuron firing rates, changes of the population average firing rate would correspond to a linear axis in the PC space (red vectors in Fig. 2I-J). By projecting a perpendicular line to this axis, one can determine the average firing rate associated with each point on the manifold. In the face task, the curved manifold is rotated with respect to the average rate axis such that the projections result in a reversed order of average firing rates for different stimulus strengths supporting the T_in_ choice. By contrast, the position of the curved manifold with respect to the average rate axis in the motion task did not create such a reversal. Thus, the apparent mismatch of average firing rates between the two tasks was incidental to the position of the manifolds, which changed substantially in the state space for the two tasks, evidenced by relative positions with respect to the average rate axis. However, the population response manifolds of both tasks shared a common geometry that orderly represented the evidence supporting each choice.

These findings indicate that LIP activity during decision formation is better characterized as a state on a curved manifold than as average firing rate. This insight leads to several important questions about the relationship between neural responses and behavior, as well as the underlying computational mechanisms. First, the curvature of the manifold suggests the possibility of an explicit code for stimulus difficulty in conjunction with the DV. Does such a code exist and if so, does the encoded task difficulty bear on the monkey’s confidence? Second, it has been widely assumed that evidence integration has a common neural representation across perceptual tasks. That is, the integrated sensory information differs across tasks, but not the representation of the integration process. Could the curved manifold for integrated evidence be fixed across tasks and what are the implications of variations across tasks? Third, is the curvature of the integration manifold compatible with existing models for network mechanisms of decision making? Finally, is the curvature of the manifold unique to LIP or is it also present in other cortical regions implicated in the decision-making process?

### Functional implications of curved manifold for evidence integration

The curved manifold indicates that LIP simultaneously encodes two key variables for the decision-making process: stimulus difficulty and integrated evidence (the DV). Stimulus difficulty, or more generally the experimentally-controlled reliability of incoming evidence, is defined as unsigned stimulus strength in both tasks and encoded along the major axis of the curved manifold (Fig. 3A). In contrast, the DV is encoded as position along the curved manifold, or alternatively as projections onto a linear axis perpendicular to the major axis of the curved manifold. While encoding of the DV has been widely observed in past experiments (Gold and Shadlen 2007; Shadlen and Kiani 2013), the majority of those experimental studies used the average firing rate as a proxy for the DV (Kumano et al. 2016; Law and Gold 2008; Mazurek et al. 2003; Shushruth et al. 2018, but see Hou et al. 2019), whereas we show that this proxy could be misleading due to the task-dependent curvature of the manifold. On the other hand, the explicit encoding of stimulus difficulty has not been reported before, to our knowledge. Rather, previous studies suggested only an implicit code for stimulus difficulty through the representation of the DV and choice (Kiani and Shadlen 2009; Pouget et al. 2016). Here we define explicit as linearly decodable (Misaki et al. 2010), and implicit as readable albeit not linearly (the explicit/implicit labels are chosen only to facilitate communication of the observed phenomenology). The curvature of the manifold, however, suggests explicit encoding of stimulus difficulty. This raises an intriguing possibility that the curved manifold contributes to both the choice and the confidence associated with the choice, as it enables linear readouts of both the DV and stimulus difficulty by downstream areas.

**Figure 3:**
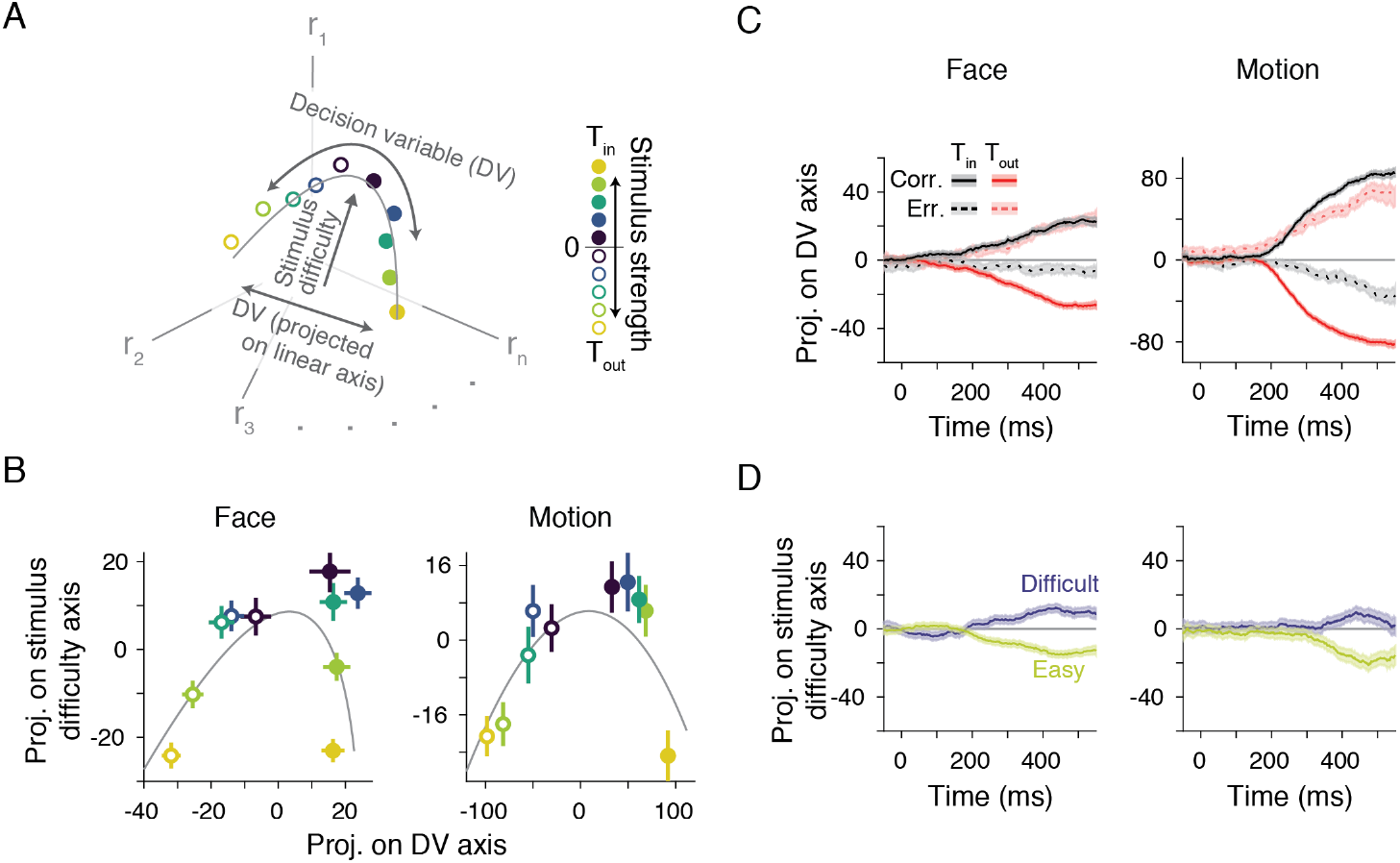
LIP population responses jointly encode the DV and stimulus difficulty. (**A**) Schematic illustration of joint encoding of the DV and stimulus difficulty with the curved manifold. The DV encoding along the curve could be approximated by a linear axis perpendicular to the major axis of the manifold. (**B**) We used an orthogonal canonical correlation analysis (CCA) to identify the best linear axes that encoded the DV and stimulus difficulty. These two axes enabled a targeted 2D projection of population responses. The plots show projected responses at 400 ms after stimulus onset (100 ms window). Similar response patterns were also observed at earlier or later times during decision formation (300-600 ms). The analysis was cross-validated; the encoding axes were identified by applying CCA to a random half of trials, and the projected data points were generated using the other half. The gray curve illustrates a manifold obtained from fitting second-order polynomials to the difficulty or DV-axis projections as a function of stimulus strength. Error bars are s.e.m. (**C**) Time course of the population neural responses projected on the DV axis for each stimulus category (T_in_ or T_out_) and for each choice outcome (correct or error). The responses were predictive of the monkey’s forthcoming choices. Note that because we combined units recorded in separate sessions, the reported values ignore the effect of neural correlations (Moreno-Bote et al. 2014). However, similar results were also obtained with simultaneously recorded units (see Fig. S5C). The shading indicates s.e.m. (**D**) Time course of the population neural responses projected on the stimulus difficulty axis for easy (≥ 20% stimulus strength) and difficult (< 20%) stimuli.

To examine this prediction, we used a three-pronged approach, where we first identified the best linear axes in the state space for reading out the DV and stimulus difficulty, then we quantified the sufficiency of these axes for predicting choice and stimulus difficulty, respectively, and finally, we tested whether projections along the stimulus difficulty axis were predictive of the monkey’s confidence. In both the motion and face tasks, the DV for a fixed stimulus duration is proportional to the signed stimulus strength (Okazawa et al. 2018; Okazawa et al. 2020; Palmer et al. 2005). Further, in both tasks, difficulty is defined by the absolute stimulus strength. We used an orthogonal canonical correlation analysis (CCA) (Cunningham and Ghahramani 2015) to find a pair of linear axes in the state space, one most correlated with the signed stimulus strength (“DV axis”) and another most correlated with the unsigned stimulus strengths (“stimulus difficulty axis”). These two axes provided a two-dimensional (2D) perspective of the population neural responses, which reproduced the curved manifold in both tasks (Fig. 3B). The weights of neurons in the two axes were unimodally distributed, with no systematic relationship between the two sets of weights (Fig. S6B), suggesting a mixed representation of the two variables and the absence of distinct functional groups in the population. The DV and difficulty encoding axes were stable throughout the decision formation period, reflecting the stability of curved manifolds in the 3D PC space (Fig. 2G-H).

Critically, the neural responses projected on the DV and difficulty axes were predictive of the choice and stimulus difficulty, respectively (Fig. 3C-D). To ensure that these predictions were not artificially introduced by our analysis, we adopted a cross-validation approach, where we first derived the DV and difficulty axes from a random half of the trials, and then used the other half for the predictions. Projections along the DV axis gradually diverged for T_in_ and T_out_ choices during stimulus viewing (Fig. 3C; *p* < 0.001 in both tasks, bootstrap test of neural responses 350–450 ms after stimulus onset). Further, when the monkey made an error, the neural responses followed a trajectory opposite to that of correct trials, indicating that the DV axis affords a representation of the DV that shapes the choice, not a representation of the objective stimulus (also see the approximate symmetry of correct and error choice in Fig. 4C). Finally, the neural responses projected on the stimulus difficulty axis did not predict the monkey’s choices (Fig. S6A), but they clearly distinguished easy and difficult stimuli (Fig. 3D; *p* < 0.001).

**Figure 4:**
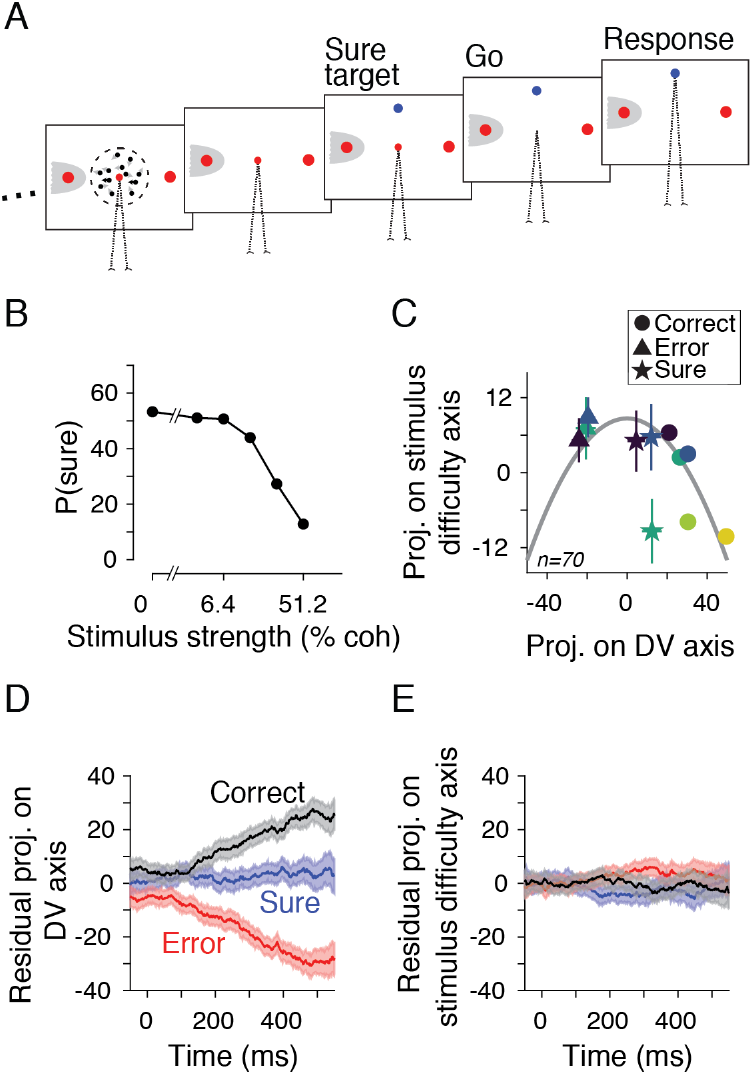
Explicit encoding of stimulus difficulty in LIP population responses is not predictive of confidence judgments. (**A**) We measured the monkey’s confidence about motion direction using a post-decision wagering task. A sure target (T_s_) appeared during the delay period on a random half of trials. The monkey could choose T_s_ to opt out of direction discrimination for a small but guaranteed reward. (**B**) Monkeys chose T_s_ more frequently for weaker motion strengths. (**C**) Population neural responses for each stimulus strength and choice 400 ms after stimulus onset (100 ms window). The sign of the DV axis is redefined with respect to the correct target such that positive DV indicates neural support for the correct choice. For cross-validation, we used a random half of the correct trials to determine the CCA axes and then projected the responses of the other half of correct trials, as well as those of the error and T_s_ trials, on these axes. The color code for stimulus strength is the same as in Fig. 3B. For error and T_s_ choices, we show only the three weakest stimulus strengths, where these choices were present in all recording sessions. Error bars are s.e.m. (**D**) Residual projected population responses on the DV axis. The residual projections were computed by subtracting the mean projection for each stimulus strength along the DV axis and then combining the residuals across different stimulus strengths. The shading indicates s.e.m. (**E**) Residual projected population responses on the stimulus difficulty axis.

Does this stimulus difficulty encoding play a role in confidence judgments? We examined this possibility by analyzing neural responses while monkeys performed post-decision wagering (Kiani and Shadlen 2009; Middlebrooks and Sommer 2012) in the motion task. In this version of the task, a third target (sure target, T_s_) appeared during the delay period on half of the trials (Fig. 4A). By choosing this target after the Go cue, the monkey could opt out of direction discrimination for a small but guaranteed reward. It has been previously verified that monkeys choose T_s_ based on their certainty, selecting it more frequently on difficult trials (Fig. 4B) (Kiani and Shadlen 2009; Komura et al. 2013; Odegaard et al. 2018). Further, previous studies demonstrated that the neural representation of the DV is predictive of the monkey’s T_s_ choices (Kiani and Shadlen 2009). However, if the neural mechanisms of confidence rely on explicit encoding of stimulus difficulty by the LIP population, projections along the difficulty axis in the state space would also be predictive of T_s_ choices. In fact, it is even possible that previous reports of the relationship between the DV representation and T_s_ choices were merely side effects of the joint encoding of the DV and stimulus difficulty in the same curved manifold, as changes of stimulus strengths across trials modify projections along both the DV and difficulty axes.

Although projections along the stimulus difficulty axis distinguished difficult and easy trials (Fig. 3D), these projections were not predictive of the monkey’s confidence (Fig. 4). To illustrate this point, figure 4C shows the projection of neural responses in the 2D space defined by the DV and difficulty axes at an example intermediate time during decision formation (400 ms after stimulus onset). For visual clarity and to better depict functional organization of neural responses, we show correct trials for all stimulus strengths, and error and T_s_ trials for the three weakest strengths where error and T_s_ choices were present in all sessions. The DV axis in this panel is thus defined with respect to the correct choice, with positive and negative values indicating the DVs in favor of the correct and error choices, respectively. Whereas different choices were associated with distinct projections along the DV axis, projections along the stimulus difficulty axis did not clearly distinguish low-confidence T_s_ choices from the higher-confidence choices where the monkey rejected T_s_ and chose one of the direction targets (correct and error choices).

To quantitatively test these observations, we asked whether, for a fixed stimulus strength, residual projections along the DV and difficulty axes were predictive of confidence over the course of decision formation (Fig. 4D, E). The residuals were computed separately for each stimulus strength and CCA axis using the mean projection across the three choices. Then, we combined the residuals of different stimulus strengths for each choice. This residual analysis addresses the stimulus difficulty confound that causes joint variations along the DV- and difficulty-axes, as it isolates the effect of variations along one axis while keeping projections on the other axis constant. The residual projections along the DV axis showed a clear separation for the three choices, with the residuals for T_s_ choices in between those for correct and error choices (Fig. 4D; correct vs. T_s_, *p* < 0.001; error vs. T_s_, *p* < 0.001, bootstrap test, 350–450 ms window). Thus, the DVs closer to zero were associated with T_s_ choices and errors occurred because the DV supported the opposite choice (Figs. 3C and 4C), consistent with previous reports (Kiani and Shadlen 2009). In contrast, residual projections along the difficulty axis failed to distinguish the three choices (Fig. 4E; correct vs. T_s_, *p* = 0.52; error vs. T_s_, *p* = 0.17), indicating that this axis did not have noticeable bearing on the monkey’s confidence. These results suggest that the functional role of the curved manifold is primarily to encode the DV.

### Distinct encoding of the DV and action plan

The curvature of the neural manifold during decision formation has important implications about the neural implementation of the decision-making process. Existing theories assume neural modules competing for their preferred actions to implement computational mechanisms of evidence integration —e.g., point attractor dynamics (Verdonck and Tuerlinckx 2014; Wong and Wang 2006), line attractor dynamics (Ganguli et al. 2008; Koulakov et al. 2002; Mante et al. 2013), or probabilistic population codes (Beck et al. 2008; Hou et al. 2019). For example, bi-stable point attractor dynamics for binary decisions have been instantiated by two action-encoding neural populations that mutually inhibit and self-excite (e.g., Wang 2002; Wong and Wang 2006; Fig. 5E). These specific implementations suggest that the DV is represented through the balance of activity of the two competing modules: the more evidence in favor of an action, the higher the responses of neurons encoding that action. However, the reversed order of T_in_ PSTHs with stimulus strength in the face task suggests otherwise. Note that LIP neurons in both tasks are selective for action, evidenced by separation of T_in_ and T_out_ PSTHs. However, the representation of the DV during decision formation is not achieved by scaling the firing rate of neurons based on the evidence supporting their favorite action.

**Figure 5:**
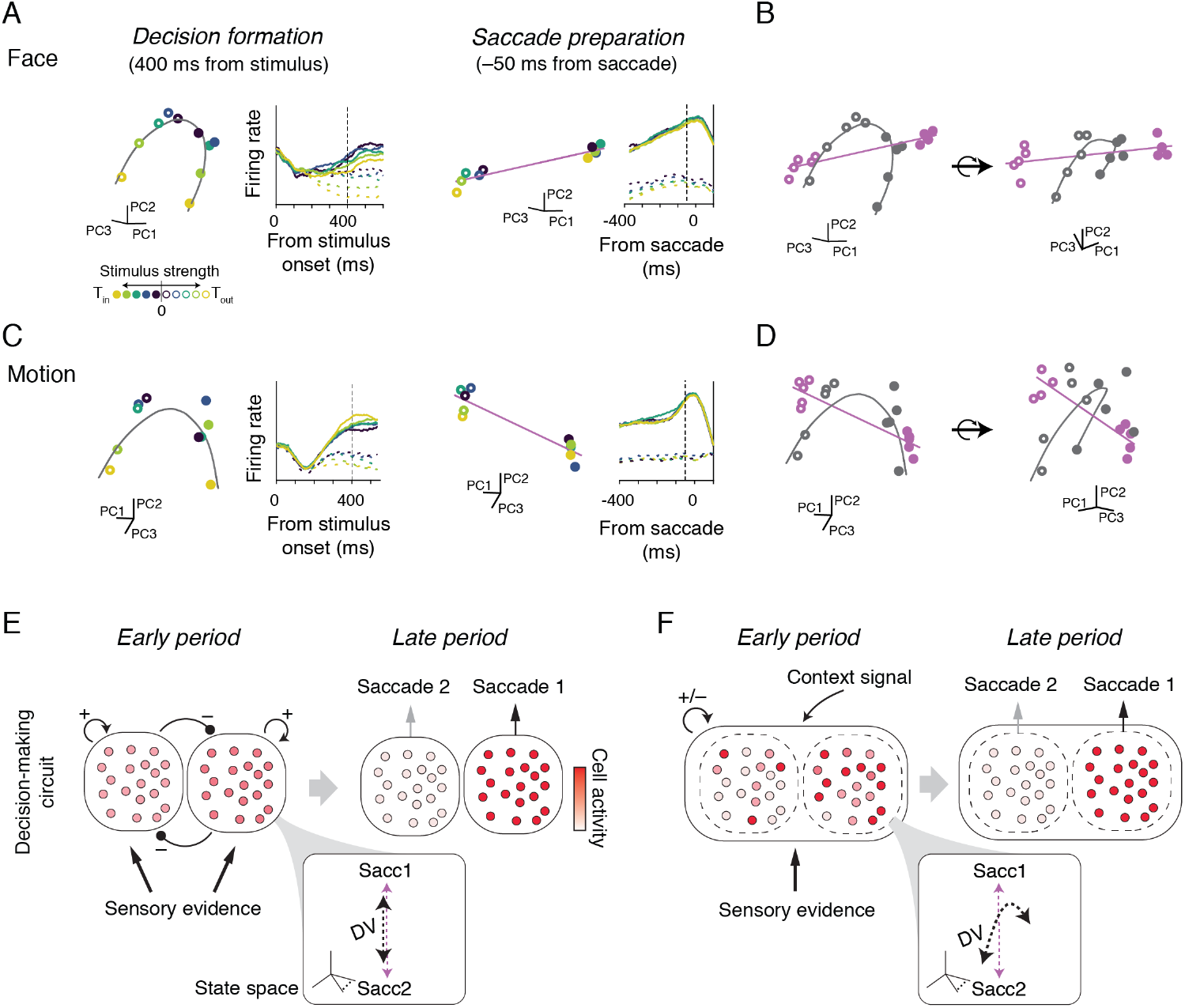
Distinct encoding of the DV and action plan. (**A**) Left: Population responses at 400 ms after stimulus onset and the average PSTHs aligned to stimulus onset in the face task (identical to Fig. 2A, G). Right: Population responses at 50 ms before saccade onset and the average PSTHs aligned to the saccade onset. (**B**) Separation of the DV and action encoding is visualized in the 3D state space by comparing the population response manifolds during decision formation (black) and saccade preparation (purple). Two different perspectives are shown to illustrate the 3D configuration. (**C-D)** Same as A-B but for the motion task. (**E**) Schematic illustration of a typical network that renders decisions through competition of groups of neurons preferring different choices or action plans (Wong and Wang 2006). Competition of action-representing neurons is commonly assumed in a wide variety of existing models to instantiate computational mechanisms of evidence integration (Beck et al. 2008; Mazurek et al. 2003; Purcell et al. 2010; Wang 2002; Wimmer et al. 2015). In these models, sensory responses supporting a specific choice increase the activity of neurons representing the associated action. These models predict a monotonic ordering of the average activity of each neural group with stimulus strength, and alignment of the DV and action encoding in the state space (inset; the dashed black line indicates the DV axis and the purple line indicates the axis connecting the states of two actions). (**F**) Alternatively, neural activity during decision formation could encode the DV in an way partly dissociated from the action selectivity of neurons (black curve in the inset is angled but not orthogonal to the action axis). As the decision-making process progresses, the DV is transformed to a choice representation through changing the activity of neurons according to their action selectivity. Consequently, there is a non-monotonic transformation of activity patterns from earlier periods reflecting the decision formation to later periods when the representation of the plan of action becomes dominant.

Although the reversed order of T_in_ PSTHs in the face task is quite revealing, the distinction of the DV and action encoding can be demonstrated in the state space of both tasks by comparing the population response manifolds during decision formation and saccade preparation (Fig. 5A-D). During saccade preparation, responses coalesced into one of two states corresponding to the two choices (Fig. 5A, C right). Correspondingly, the average firing rates attained a common level for each choice (Fig. S4). However, the curved manifold of the DV during decision formation was misaligned with the axis connecting the two action states (Fig. 5B, D). The angle between the two axes (face task, 34.4°±4.0°; motion task, 11.7°±1.6° in 3D space) was significantly greater than the expected angles based on the variability of neural responses (*p* < 0.001, bootstrap test). The expected angles based on response variability were estimated by comparing the choice axes between random halves of trials (face task: 2.9°±1.6°, motion task: 2.5°±1.3°). The misalignment with the states encoding the two actions indicates that the relative activity of two pools of action selective neurons does not provide a veridical representation of the DV during decision formation (see also Figs. 2E and S2D).

This form of the DV encoding does not naturally arise in existing network models, in which competition among neuronal modules associated with different action plans underlies decision formation (Fig. 5E; Beck et al. 2008; Deco et al. 2013; Mazurek et al. 2003; Purcell et al. 2010; Wang 2002; Wimmer et al. 2015). Therefore the curvature of the neural manifolds and the dissociation from the action encoding should originate from a mechanism not considered in previous models. Our experimental data suggest that such a mechanism should realize a non-monotonic transformation of the neural activity from the DV encoding in early phases of decision formation to encoding of a planned action at later times in the trial (Fig. 5F). In other words, the encoding of the DV in LIP is not tightly bound to the actions associated with the decisions, especially in early stimulus viewing periods. At later times, however, the DV representation gives rise to the representation of the upcoming saccade and is ultimately supplanted by it.

### Task dependency of the DV encoding

Another common assumption in existing models is that different perceptual tasks that rely on integration of evidence are distinguished by only the information that is being accumulated; the neural representation of the integration process is presumed largely identical across tasks. LIP neurons had dissimilar responses in the face and motion tasks (Fig. 2). But we could not rule out that these response dissimilarities stemmed solely from differences of sensory inputs to the LIP (motion vs. face). Could task demands be a major factor? Specifically, would the DV encoding change for different decisions made with the same sensory stimuli? To answer this question, we analyzed a subset of neurons (n = 28) recorded in both the species and expression categorizations in the face task. Recall that the face stimuli in each session could vary along two morph axes: species and expression. The stimuli around the center of this 2D space were presented in both categorization tasks (purple box in Fig. 6A). Thus, monkeys applied two different categorization rules to the same visual stimuli.

**Figure 6:**
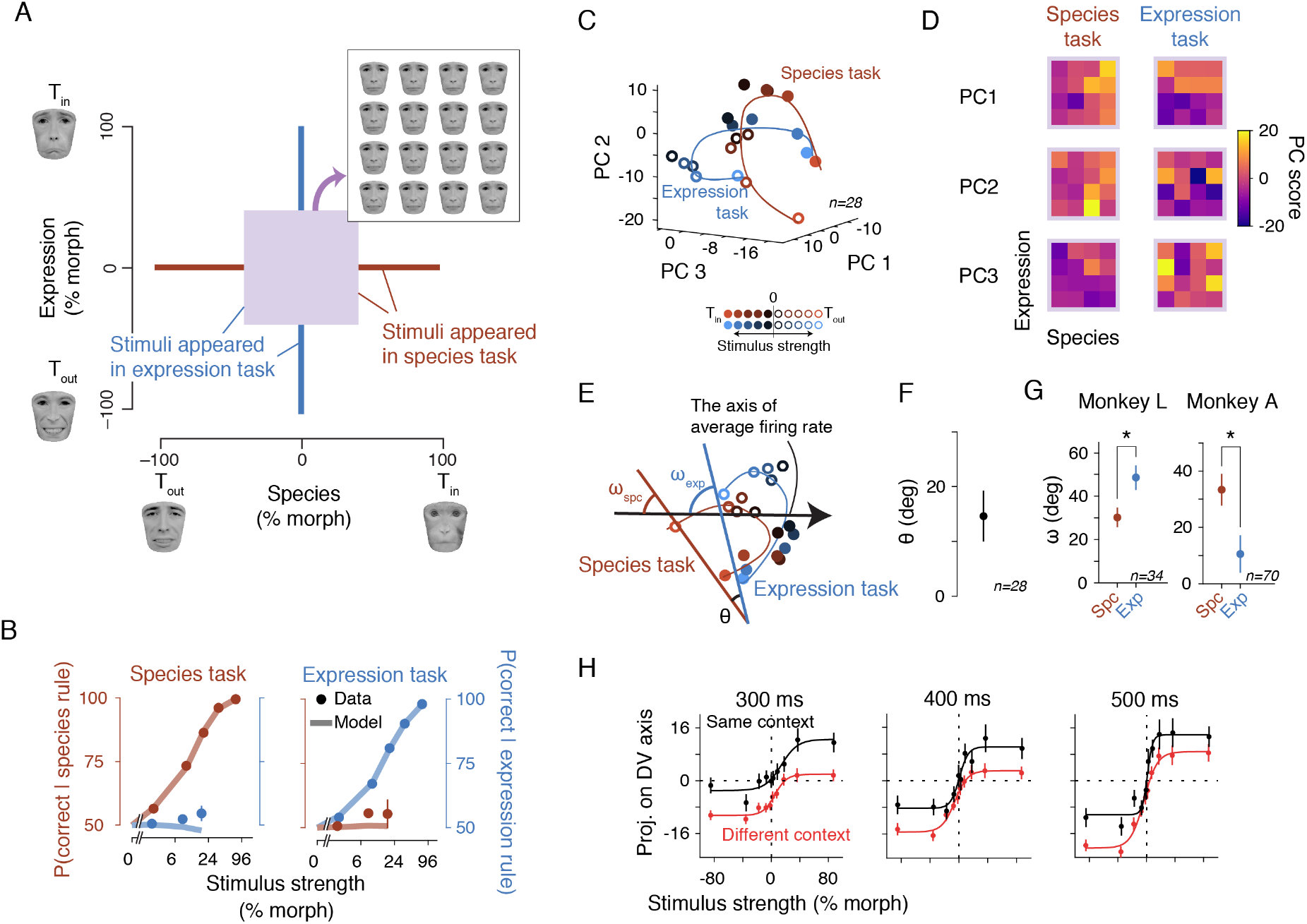
Task dependency of the population response manifold. (**A**) We examined task dependency of LIP responses by comparing the species and expression categorizations. The face stimuli used in these tasks can be visualized in a 2D space encompassing the species and expression morphs. The stimuli at the central region of this space (purple box) appeared in both tasks. (**B**) Performance in both task contexts with respect to each categorization rule (red: species, blue: expression). Near chance performance for the irrelevant categorization rule (e.g., blue dots in the species task) indicates that monkeys correctly ignored the irrelevant stimulus fluctuations. Lines are drift diffusion model fits. Error bars are s.e.m. (**C**) The neural manifolds were distinct in the two task contexts. The PCs were derived from the neural responses across both task contexts. The manifolds were generated by projecting population responses of each task onto the common PC axes. The plot was generated using a subset of units (n = 28) recorded in both tasks. (**D**) The same stimuli elicited different response patterns. The heat maps show projections of population responses on the top three PCs for the common stimuli in the central region of the morph plane (purple box in A). We divided this region into 16 sub-regions for better visualization. The relevant category boundaries are apparent in the first PC projections. (**E**) The same manifolds in C from a different perspective to allow a better illustration of angles between the DV-encoding axes of the two manifolds (*θ*), as well as their angles with the average rate axis (*ω_spc_* and *ω_exp_*). The DV-encoding axes (straight lines running parallel to the curved manifolds) were computed using CCA. (**F**) *θ* in the 3D state space was significantly greater than zero. (**G**) *ω_spc_* and *ω_exp_* were significantly different in both monkeys. Unlike *θ* which could be estimated only with units recorded in both tasks, the calculation of *ω*’s could be done using all units. Here we show results from units recorded in only one task so that F and G provide independent support for our conclusion. (**H**) Using the DV-encoding axis of one task to read out the DVs in the other task caused substantial biases. The plots show neural responses of expression categorization task projected on the DV encoding axis of the same task or the species categorization task. Projecting on the DV axis of the species task leads to a strong bias absent in behavior. Similar biases were present when the DV axis of the expression categorization task was used for decoding neural responses of the species categorization task. The plots were cross-validated to ensure that the trials used for deriving the DV axes did not overlap with those used for creating the projections. The fitting curves are logistic functions (Eq. 9).

Monkeys successfully performed both categorization tasks without significant interference from the orthogonal task rule (Fig. 6B). In both contexts, PSTHs showed a reverse order with stimulus strength (Fig. S2C), and there was significant curvature in the population response manifolds (Fig. 6C, *p* < 0.001). Critically, the manifolds formed by the neural activity within the species and expression categorization contexts were quite distinct from each other (Fig. 6C; different coefficients of polynomial fits to the manifolds, *p* = 2.8 × 10^−15^, likelihood-ratio test). These manifold differences were apparent in population response patterns for the same stimuli in the two tasks, evident in the projections of population responses on the top three PCs (Fig. 6D; *p* = 0.012, permutation test). Projections on the first PC were most striking, as they revealed the distinct category boundaries of the two task contexts (top row of Fig. 6D, a vertical boundary in the species task and a horizontal boundary in the expression task).

The differences in the manifolds caused the linear DV axes derived from CCA to be different in the two task contexts. The two DV axes projected onto the 3D state space had a +15.1° ± 5.7°, angle (signed with respect to the axis of average firing rate), which was significantly different from 0° (Fig. 6E-F; *p* < 0.001, bootstrap test; the angle difference grew larger in higher dimensional state spaces). Consequently, when a DV decoder optimized for one categorization task was used for decoding the DV in the other task, a substantial readout bias emerged (Fig. 6H; Δ*γ*_0_ = 5.9 ± 1.1 in Eq. 9 for decoding with the same- and different-context DV axes at 400 ms, *p* = 3.4 × 10^−9^, likelihood-ratio test).

Although the neurons recorded in both face categorization tasks provided the best test for changes of manifolds with task context, the difference could also be inferred from the units recorded only in one categorization context (n = 104). For those units, we computed the angle between the DV axis and the average firing rate axis for each context (*ω_spc_* and *ω_exp_* in Fig. 6E). If an area encodes the DV in a context-independent manner, these angles would be identical. However, we found them to be significantly different (Fig. 6G; Monkey A, *p* = 0.006; Monkey L, *p* = 0.006, bootstrap test). Together, the manifolds that encode the DV changed in the state space depending on task contexts, and the observed differences could not be attributed solely to the differences of sensory stimuli.

### The DV encoding in lateral and medial prefrontal areas

Our analyses thus far focused on LIP neural activity. However, decision-related signals encoding the DV are prevalent across multiple brain structures (de Lafuente et al. 2015; Deverett et al. 2018; Ding and Gold 2010, 2012; Horwitz and Newsome 1999; Kiani et al. 2014b; Kim and Shadlen 1999). If the curved manifold is a representational property of decision formation, areas other than LIP are also expected to have this response geometry. Alternatively, the manifold curvature could be specific to LIP and stem from its circuit and functional properties. To test these possibilities, we examined the responses of two other cortical areas in the frontoparietal network involved in oculomotor decisions during the motion task: the supplementary eye field (SEF) and the prearcuate gyrus (PAG) (Fig. 7).

**Figure 7:**
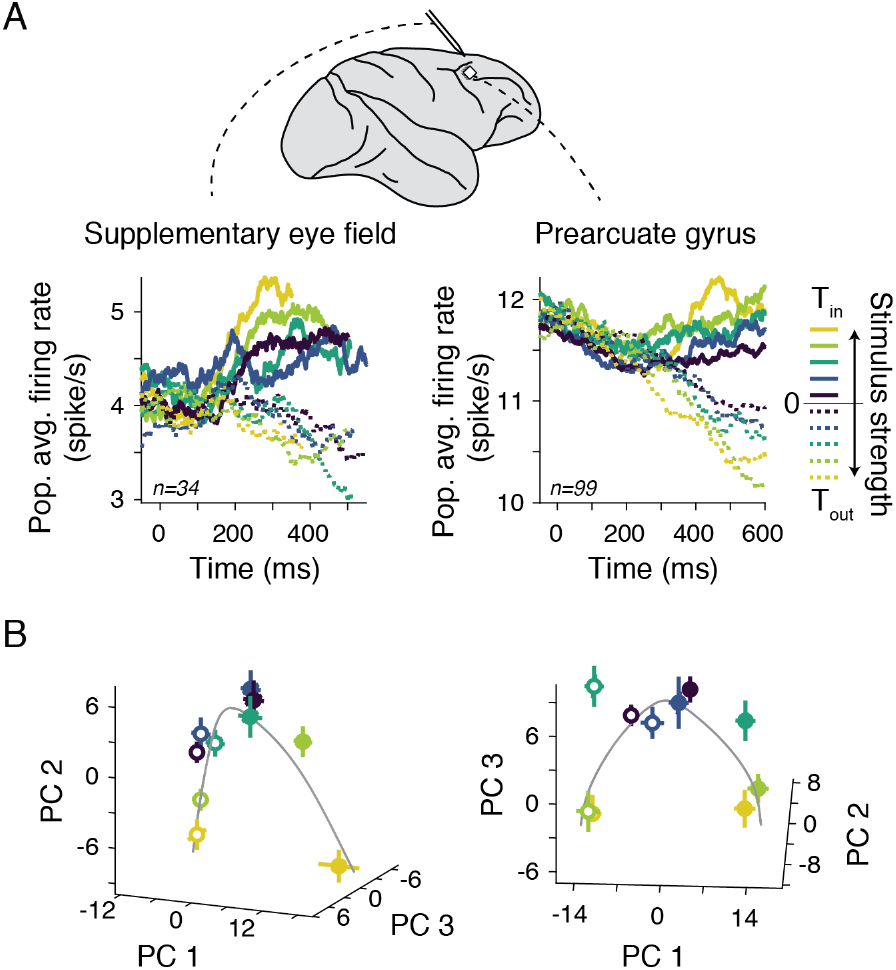
Curved population response manifolds in the lateral and medial prefrontal cortices. (**A**) PSTHs of prearcuate gyrus (PAG) and supplementary eye field (SEF) neurons during a reaction time version of the motion task. Both areas showed monotonically increasing responses for larger stimulus strengths during T_in_ choices, similar to LIP. PAG and SEF were recorded with Utah arrays and linear probes, respectively, shown on the lateral view of the macaque brain (top panel). (**B**) Population responses in both areas formed a curved manifold as in LIP. The plots show the manifold at 400 ms after stimulus onset (100 ms window). Conventions are the same as in Fig. 2G-H.

Both areas revealed similar response geometries as in LIP. Critically, population responses, visualized using the same PCA procedure as for LIP, formed a curved manifold in both areas (Fig. 7B; significance of curvature, *p* < 0.001), with a monotonic arrangement of different stimulus strengths along the manifold. In these recordings, monkeys performed a reaction time variant of the task, where they indicated their decision as soon as they were ready. In the reaction-time task, the monkey did not wait for a Go cue and the trial terminated as soon as the monkey committed to a choice. Hence the results confirmed that the curvature of the manifold is indeed a property pertaining to decision formation, not a post-commitment representation and not a persistent representation of choice. Together, the results across the frontoparietal network indicate that the curvature of the neural manifold that encodes the DV is a ubiquitous representational property for the brain regions involved in decision formation.

## Discussion

Brain regions involved in motor planning also represent formation of decisions conveyed through actions (Cisek and Kalaska 2005; de Lafuente et al. 2015; Ding and Gold 2012; Donner et al. 2009; Horwitz and Newsome 1999; Kim and Basso 2008; Ratcliff et al. 2003). For example, in a direction discrimination task with random dots, the average firing rate of saccade-selective LIP and FEF neurons increases with larger evidence supporting the preferred saccade direction of the neurons (Ding and Gold 2012; Kim and Shadlen 1999; Mante et al. 2013; Shadlen and Newsome 2001). This encoding of the decision variable (DV) is observed only when decisions are associated with specific actions (Bennur and Gold 2011; Wang et al. 2019), and when such associations exist, neurons appear to encode the DV as a partially enabled (potentiated) action plan (Cisek and Kalaska 2010; Shadlen et al. 2008). Inspired by these experimental observations, existing theoretical and circuit models of the decision-making process explain decision formation as a competition of neurons representing potential actions (Beck et al. 2008; Mazurek et al. 2003; Purcell et al. 2010; Wang 2002; Wimmer et al. 2015). Furthermore, because decisions are achieved through competition of actions, these models also assume that the DVs of different decisions share the same neural code (Hou et al. 2019) (and therefore a common decoding scheme (Mante et al. 2013)) as long as they involve the same competing actions, regardless of the sensory or contextual inputs that differentiate the decisions. Our results challenge both of these fundamental assumptions.

We show that although decision formation is represented in the activity of neurons that ultimately encode actions (de Lafuente et al. 2015; Dorris and Glimcher 2004; Shadlen and Newsome 2001; Snyder et al. 2000), the neural responses during decision formation do not necessarily scale with the evidence supporting the preferred action of the neurons, even when the decision is communicated through that action. This is best observed in our face task, where PSTHs show a reverse order: decreased firing rates with stronger supporting evidence for T_in_ choices (Fig. 2A). Instead of scaling neural responses by the evidence supporting the preferred action, we find a curved linear manifold in the state space of the neural population that represents the DV (Fig. 2G-H). Depending on the rotation of this manifold with respect to the axis of average firing rate, population PSTHs may show an arbitrary, non-monotonic order with evidence (Fig. 2I-J). Our results highlight the conceptual gap between average firing rates and the DV (Raposo et al. 2014; Schall 2019), as well as clarify the form and geometry of the neural code for the decision-making process.

The task-dependent changes of the DV manifold are not simply due to changes in sensory stimuli across tasks. They occur even when the stimuli are kept constant but task rules are changed (Fig. 6), suggesting that contextual factors shape the neural computations beyond what is assumed by existing models. Population level analysis is key to elucidating and understanding context-dependence of the decision-making process, as such dependencies could be obscured in average firing rates (Hou et al. 2019; Kumano et al. 2016). Critically, the rotation of DV manifolds implies that accurate readout of the DVs —a necessary process for action planning and circuit-circuit coordination in the decision-making network— must take task context into account. In the absence of such “context-aware readouts,” there would be substantial inaccuracies manifesting as choice bias (Fig. 6H).

Why does the curvature exist ubiquitously in all the tasks and brain regions studied here? We suggest that curved manifolds arise automatically due to fundamental constraints on neural computations (Fig. S7). The first set of constraints limit the dynamic range of the firing rates (and hence projections along the encoding axes) that constitute the neural code: the non-negativity of firing rates puts a lower bound on the dynamic range, and various costs associated with spiking (e.g., metabolic costs, Lennie 2003) curtail firing rates from above. An energetically efficient code would seek to lower the overall firing rates of neurons, bending the manifold near the non-negativity bound (Keemink and Machens 2019; Fig. S7C). The second set of constraints arise from the need for a precise code. Encoding a continuous decision variable within a limited dynamic range could become more precise with a curved manifold than a straight line, as the difference of nearby DV values could be encoded with larger, more distinguishable changes in population responses (Fig. S7C). However, the exact precision gained by a curved manifold depends on the correlation structure of the population (Bondy et al. 2018; Moreno-Bote et al. 2014) and the feasibility of decoding along the curved manifold instead of a linear readout. Fortunately, decoding along a low-order polynomial in state space, similar to those in our study, is not prohibitively challenging. Further, there is no strong reason to favor a linear readout over other simple readouts (Ritchie et al. 2019). However, the complexity of the manifold cannot increase in an unbounded fashion as the resulting neural code may become susceptible to decoding errors or become too tangled to be easily readable (Fig. S7E). We propose that for each task, the sensory and contextual inputs define the sub-region of the state space where the DV can be encoded in, and the constraints above shape a curved manifold that accommodates adequately precise encoding of the DV and an easy readout.

The curved manifold indicates that the LIP population activity explicitly encodes both stimulus difficulty (i.e., reliability of sensory evidence) and the DV (Fig. 3A), a departure from past studies that reported only implicit encoding of stimulus difficulty by single neuron responses in LIP (Kiani and Shadlen 2009; Pouget et al. 2016). This encoding of stimulus difficulty cannot be explained away as an increased attention level or task engagement for more difficult stimuli. Two key observations corroborate our conclusion. First, stimulus difficulty is encoded from the very beginning of the representation of the decision-making process (∼200 ms after stimulus onset; Fig. 3D), making it unlikely to be reactive recruitment of engagement mechanisms due to perceived stimulus difficulty. We remind that stimulus difficulty is not predictable prior to the stimulus onset and inferring the level of difficulty from a stimulus is time-consuming (Khalvati et al. 2020). Second, and more critically, higher engagement should lead to better behavioral performance, but the activity along the difficulty encoding axis was uncorrelated with the monkey’s accuracy (Fig. S6A). Moreover, trial-to-trial variations of the encoded difficulty for the same stimulus strength did not correlate with the monkey’s confidence in a post-decision wagering task (Fig. 4), providing a clear example that decodability does not necessarily amount to functionality (Ritchie et al. 2019). This apparently non-functional encoding of difficulty lends support to the hypothesis that the manifold curvature is a geometric property of population responses for computing or monitoring changes of the DV, as explained above.

We further rule out that the curvature of the DV manifold and encoding of stimulus difficulty arise from sensory responses or spatial attention to stimuli. Parietal neurons respond to a variety of stimulus attributes in their RFs (Bisley et al. 2004; Janssen et al. 2008; Lehky and Sereno 2007; Sarma et al. 2016) and also show strong response modulations depending on the location of spatial attention (Bisley and Goldberg 2003). However, these response properties do not explain the reversal of firing rates. First, our recording from face-selective regions in the inferior temporal (IT) cortex during the face tasks has revealed that sensory responses are monotonically modulated by the stimulus strength (Fig. S2F), just like the monotonic tuning to random dots stimuli in the middle temporal (MT) area (Britten et al. 1993). Therefore, the reversal was not inherited from response properties in sensory areas. Second, our sensory stimuli were presented outside the LIP RFs. Increased sensory responses or spatial attention to more difficult stimuli would not affect or rather decrease the activity of the recorded LIP neurons, leading to a response pattern opposite to the reversed order in the face task. To further ascertain that the observed LIP responses were not affected by inadvertent overlaps of the stimuli with RFs, we performed a control experiment in our face task, where we showed the stimuli in the visual hemifield opposite to the LIP RFs. The results replicated the same reversal of firing rates and curved manifolds as in the main task (Fig. S8), rejecting the influences of sensory responses and spatial attention. It should also be noted that past studies have reported similar LIP response patterns regardless of whether motion stimuli were presented foveally or parafoveally (Huk and Shadlen 2005; Shushruth et al. 2018), indicating that differences in stimulus position across tasks do not explain our results.

Our finding that similar response geometries and encoding dynamics are present in LIP as well as lateral and medial prefrontal cortices (Fig. 7) suggests ubiquitous principles that shape the neural code throughout the cortical nodes of the decision-making network in the primate brain. Our conclusions, therefore, transcend present debates about the causal role of LIP in perceptual decisions (Jeurissen et al. 2019; Katz et al. 2016; Shushruth et al. 2018; Zhou and Freedman 2019). The cited studies have yielded diverse and occasionally incompatible results, suggesting that decisions may be formed through interactions of multiple brain regions, instead of resulting from a single region (e.g., LIP). Such a distributed computation creates some degree of robustness against local perturbations (Li et al. 2016). As we have shown here, multiple nodes within the network involved in saccade planning exhibit similar response geometries adding further credence to a multi-region network model for decision making. Pertinently, neural responses in the superior colliculus, caudate, and frontal eye fields have been occasionally reported to vary inversely or non-monotonically with stimulus strength (Ding and Gold 2010, 2012; Horwitz and Newsome 2001), hinting at the potential presence of curved DV manifolds in those regions, which would extend our discovery to the subcortical nodes of the decision-making network as well.

The geometric properties of the manifold further suggest that the DV encoding is distinct from the patterns of neural activity that represent actions. Unlike psychological models (e.g., DDM) that are agnostic to the neural implementation of the decision-making process, circuit models have to clarify the true nature of the DV, its dynamics, and its transformation to action plans. To account for our results, computational mechanisms that underlie integration of sensory evidence and commitment (termination) rules for perceptual decisions —e.g., point attractor dynamics (Verdonck and Tuerlinckx 2014; Wong and Wang 2006), line attractor dynamics (Ganguli et al. 2008; Koulakov et al. 2002; Mante et al. 2013), or probabilistic population codes (Beck et al. 2008; Hou et al. 2019)— must be implemented to untangle from encoding actions. This, however, does not indicate that decision formation takes place in circuits separated from action planning. Rather, the same LIP neurons that initially encode the DV on a curved manifold gradually change their response patterns to encode actions later in the trial (Figs. 5 and S4). This sequential use of the same neural population for decision-making and action planning matches prior experimental observations in the premotor and motor cortex (Kaufman et al. 2015; Peixoto et al. 2018), especially progressive recruitment of choice representation (Peixoto et al. 2018). Thus, the dissociation between the DV and action planning necessitates network models in which the same neural circuits undergo a fundamental transformation of population response patterns over time.

A hallmark of the decision-making process is its flexibility. To implement flexible decisions, neural circuits must accommodate context-dependent interpretation of information, intricate rules, and a large variety of motor actions to interact with the outside world. The curvature and context-dependency of the DV manifolds and partial separation of the DV and action encoding, as we report here, are some of the building blocks that the primate brain may have devised for flexible and accurate decisions based on sensory information.

## Methods

We recorded responses of parietal and frontal neurons while seven macaque monkeys (macaca mulatta) performed a variety of perceptual decision-making tasks. Two monkeys (A and L) performed two variants of a novel face discrimination task and five monkeys (D, I, S, O, and N) performed direction discrimination tasks with random dots (Kiani et al. 2008; Roitman and Shadlen 2002). Monkeys D and I also performed the dots task with post-decision wagering (Kiani and Shadlen 2009). All experimental procedures conformed to the National Institutes of Health *Guide for the Care and Use of Laboratory Animals* and were approved by the institutional animal care and use committee at New York University. The data from monkeys D, I, and S were previously published (Kiani et al. 2008; Kiani and Shadlen 2009).

### Behavioral tasks

Monkeys were seated in a semi-dark room in front of a cathode ray tube monitor (frame rate, 75 Hz) with their heads stabilized using a surgically implanted head post. Stimulus presentation was controlled with Psychophysics Toolbox (Brainard 1997) and Matlab. Eye movements were monitored using a high-speed infrared camera (Eyelink, SR-Research, Ontario) or a scleral eye coil (Fuchs and Robinson 1966). Gaze positions were recorded at 1 kHz.

Throughout the paper, we refer to the face discrimination task as the “face task,” and to the direction discrimination task with random dots as the “motion task.”

### Face task

The task required classification of faces into two categories, each defined by a prototype face. Each trial began when the monkey fixated on a small fixation point at the center of the screen (diameter, 0.3°). Shortly afterward, two red targets appeared on opposite sides of the screen, equidistant from the fixation point. After a variable delay (truncated exponential distribution, range, 250–500 ms), a face stimulus appeared 1.6°–2.5° to one side of the fixation point (contralateral to the recording site in the main experiment or ipsilateral in a control experiment, Fig. S8). The stimulus size randomly varied across trials by one octave to prevent the monkey from relying on local features (width range, 2.9°-5.8°, mean, 4.2°). The stimulus was presented for a variable duration (truncated exponential distribution, range, 227–1080 ms, mean, 440 ms) followed by a delay period (truncated exponential distribution, range, 300–900 ms, mean, 600 ms). After the delay, the fixation point disappeared (Go cue), and the monkey reported the face category with a saccadic eye movement to one of the targets. Correct responses were rewarded with a drop of juice. To manipulate task difficulty, we created a morph continuum between the two prototypes and presented different intermediate faces (see below). When the face was halfway between the two prototypes on the morph continuum, the reward was given randomly.

We trained monkeys to categorize the same face stimuli in two distinct ways: species and expression categorization (Fig. 6A). In the species categorization task, the two prototype stimuli were human and monkey faces, whereas in the expression categorization task, they were faces with “happy” and “sad” expressions. Here, happy and sad are used as designations that facilitate explaining the task. We focus merely on the monkey’s ability to discriminate different expressions and do not imply that they interpret these expressions as humans do. Monkeys performed these two categorizations in blocks that switched either within or across sessions. In the first few trials of each block, we presented only the faces closest to the prototypes of the discriminated categories to cue the task context to the monkey. In both categorization tasks, monkeys easily generalized across different facial identities. We therefore created multiple stimulus sets from different human and monkey identities. Each set enabled testing both categorization tasks (see below). We switched the stimulus set after runs of sessions. In total, three stimulus sets were used during electrophysiological recording sessions.

Each stimulus set can be visualized as a two-dimensional (2D) “face space,” whose axes correspond to the morph level between species and expression category prototypes (Fig. 6A). This 2D face space was created with three “seed” faces: photographs of happy and sad expressions of a human face and neutral expression of a monkey face. The seed faces were obtained from the MacBrain Face Stimulus Set (Tottenham et al. 2009) (http://www.macbrain.org/resources.htm) and the PrimFace database (http://visiome.neuroinf.jp/primface). We manually defined 96–118 anchor points on each seed face and developed a custom algorithm to morph the three seed faces by computing a linear weighted sum of the positions of the anchor points and textures inside the tessellated triangles defined by the anchor points. Our algorithm allowed independent morphing of different stimulus regions. Further, because our morphing changed both the geometry and content of the facial regions, the resulting faces were perceptually seamless.

Natural faces are complex and high-dimensional stimuli that challenge investigation of the decision-making process. To control stimulus complexity and make studying the decision-making process tractable, we limited informative features for face categorization. We selected three features in each face (eyes, nose, and mouth) and limited morphing across faces to those regions, reducing the effective stimulus dimensionality. Regions outside the informative features were always fixed at the midpoint of the morph space between the three seed faces and did not change in the experiment.

Each informative facial feature varied in a 2D “feature space” defined by the corresponding features of the three seed faces. Morphed features were generated as weighted combinations of the three seed features. To change the features along the species axis, the weights for the happy (*W_h_*), sad (*W_s_*), and monkey (*W_m_*) faces varied from [0.5, 0.5, 0] to [0, 0, 1]. We label these two end points as −100% and +100% species morph levels (or species prototypes). For the expression axis, the weights for the prototypes changed from [0.75, −0.25, 0.5] to [−0.25, 0.75, 0.5]. The negative weights indicate linear extrapolation beyond the seed features. We verified that for the small weights used in our stimulus sets, all morphed features and resulting faces looked naturalistic and did not show noticeable aliasing. In the 2D space, the species morph level of a feature was defined by (*W_m_* − *W_h_* − *W_s_*) × 100%, and its expression morph level was defined by (*W_s_* − *W_h_*) × 100%. Because the three informative features were morphed independently in their 2D feature spaces, our full stimulus space was six dimensional. The 2D face space of Fig. 6A shows faces with identical morph levels of eyes, nose, and mouth — a 2D cut of the 6D space.

On each trial, we chose a nominal morph level along the relevant axis for the categorization task (species axis for species categorization or expression axis for expression categorization). This nominal value determined the morph level of the three features in the trial, as well as the response that the monkey was rewarded for. The nominal morph level ranged from –96% to +96%. The exact set of nominal morph levels was adjusted for each stimulus set to keep the monkey’s overall accuracy roughly constant across the sets.

To investigate how monkeys weighted evidence conferred by the three informative features, we allowed the features to fluctuate randomly around the nominal morph level of the trial every 106.7 ms during stimulus presentation. For all nominal morph levels between −12% and +12%, the three features fluctuated independently within their own 2D feature spaces according to a circular Gaussian distribution with a standard deviation of 20% morph level. The features therefore changed both along the relevant and irrelevant axes for the categorization task, allowing us to gauge which ones influenced the monkey’s choices. For strong nominal morph levels (>12%), fluctuations happened only along the relevant axis in order to prevent the monkey from confusing the task context. Sampled values that fell outside the prototype range [–100% +100%] were replaced with new samples inside the range (5.2% of samples).

We used a masking procedure to keep changes of features in a trial subliminal (Okazawa et al. 2018). The masks were created by phase randomization of faces (Heekeren et al. 2004) and interleaved the stimulus fluctuations. In each 106.7 ms sample-mask cycle, a face stimulus was shown for 13.3 ms and then gradually faded out as the mask faded in. In the fading period, the mask and the stimulus were linearly combined, pixel-by-pixel, according to a half-cosine weighting function, such that in the last frame, the weight of the mask was one and the weight of the face was zero. In the next cycle, a new face stimulus with slightly altered informative features was shown, followed by fading with another mask, and so on. Before the first sample-mask cycle in the trial, we showed a mask frame to ensure all stimulus samples were preceded and followed by masks. The masking procedure was quite effective in pilot experiments with humans, preventing the detection of the feature changes. The stimulus in each trial looked like a face appearing and disappearing behind cloud-like patterns. Stimulus viewing duration in each trial consisted of 2-10 stimulus-mask cycles (truncated exponential distribution, mean, 4), which corresponded to 227–1080 ms, including the initial mask.

Two monkeys performed the task in 72 recording sessions (monkey A, 40 sessions; monkey L, 32 sessions), which amounted to 56,582 trials (monkey A, 18,080 and 9,309 trials for species and expression categorizations, respectively; monkey L, 17,077 and 12,116 trials). In the majority of these sessions (55), the face stimulus was contralateral to the recording site. In the control condition where the stimulus was ipsilateral to the recording site (Fig. S8), we ran 17 sessions (monkey A, 10; monkey L, 7) and collected 12,944 trials (monkey A, 7,584 trials for species categorization; monkey L, 4,041 and 1,319 trials for species and expression categorization, respectively).

### Motion task

The experimental settings for the motion task are described elsewhere (Kiani et al. 2008; Kiani and Shadlen 2009), and summarized here. The monkey began each trial by fixating a fixation point at the center of the screen, followed by the appearance of two red targets on opposite sides of the screen equidistant from the fixation point. After a variable delay (truncated exponential distribution, range, 250–600 ms), the moving random-dots stimulus (Britten et al. 1992) appeared within a 5° circular aperture centered on the fixation point. The percentage of coherently moving dots determined the strength of motion (coherence). The motion strength varied randomly across trials according to a uniform distribution from the following set: 0%, 1.6%, 3.2%, 6.4%, 12.8%, 25.6%, and 51.2% coherence. The net motion direction was toward one of the two targets and varied randomly across trials. Monkeys I and S performed a variable duration task, where the stimulus was presented for 80-1500 ms (truncated exponential distribution, mean, 311 ms). The Go cue occurred after the stimulus or following a delay period (truncated exponential distribution, range, 500–1000 ms, mean, 847 ms), instructing monkeys to report their perceived motion direction with a saccadic eye movement to one of the targets. Monkey D as well as monkey I performed another variable duration task, where the stimulus duration was 100-900 ms (mean, 286 ms) and always followed by a delay duration of 1200-1800 ms (mean, 1386 ms). Monkeys O and N performed a reaction time task, where they were free to respond any time after the stimulus onset (mean reaction time, monkey O, 729 ms, monkey N, 927 ms). Correct responses were rewarded with water or juice. For 0% coherence motion, the reward was given randomly.

Monkeys D and I also performed a version of the variable duration task with post-decision wagering (Kiani and Shadlen 2009). During this task, on a random half of the trials, a third target (“sure” target) appeared at a random time during the delay period (500–750 ms after motion offset) and stayed on through the rest of the delay period. After the Go cue, the monkey made a saccade either to one of the two direction targets or to the sure target, if present. Choosing the sure target always yielded a reward, but the reward size was smaller than that for choosing the correct direction target. The reward ratio was adjusted to encourage the monkey to choose the sure target on nearly half of trials.

We collected 67,349 trials in the variable duration task (monkey I, 24,603 trials; monkey S, 21,947 trials; monkey D, 20,799 trials). Among them, 30,991 trials were in the post-decision wagering task (monkey I, 10,192 trials; monkey D, 20,799 trials). We collected 23,756 trials in the reaction time task (monkey O, 6,291; monkey N, 17,465). Psychophysical thresholds were comparable across the tasks.

### Neural recording

For recordings from the lateral intra-parietal (LIP) cortex, electrodes were placed in the ventral division of LIP (LIPv), located based on both structural magnetic resonance imaging scans and transition of white and gray matter during recordings. The recording was performed through a plastic cylinder (Crist Instruments, Damascus, MD), implanted on the skull, and a plastic grid (1 mm spacing; Crist Instruments), placed inside the chamber for precise targeting of the electrodes. We used either single tungsten microelectrodes (FHC, Bowdoin, ME) or 16-channel linear array probes (V-Probe; Plexon, Dallas, TX). Action potential waveforms were isolated online using a time-amplitude window discriminator or sorted offline (Plexon offline sorter). In our analyses, we combined both well-isolated single units and multi-units. We confirmed that similar results were obtained using single units alone (Fig. S5D).

Units were selected for analysis if they exhibited spatially-selective persistent activity during the delay period of a memory-guided saccade task (Gnadt and Andersen 1988) (Fig. S9A-B). While the monkey maintained fixation, a target briefly flashed on the screen and was followed by a delay period (∼1000 ms). At the end of the delay period, the fixation point turned off (Go cue), and the monkey made a saccade to the remembered target location. The target location varied randomly across trials. The response field (RF) of the neurons was identified as the target locations associated with the largest firing rates during the delay period. During the motion or face task, we placed one of the targets (T_in_) in the RF and the other (T_out_) outside the RF. For the post-decision wagering results included in this paper, the sure target (T_s_) was placed outside the RF.

Our choice to focus on units with persistent delay activity in a memory-guided saccade task was made to remain consistent with previous studies (Roitman and Shadlen 2002; Shadlen and Newsome 2001). However, the LIP neurons lacking persistent activity also showed similar curved population response manifolds during our main tasks (Fig. S9C-F). Thus, similar conclusions could be made using all LIP neurons or different subgroups of neurons.

In the motion task, 129 units were recorded from the LIP of three monkeys (monkey D, 46 single units; monkey I, 50 single units; monkey S, 33 single units). In the face task, 132 units were recorded from two monkeys (monkey A, 70 (single: 41); monkey L, 62 (single: 34)) with face stimuli presented contralateral to the recording site. An additional 44 units (monkey A, 31; monkey L, 13) were also recorded during the control condition with ipsilateral stimuli (Fig. S8).

To examine the decision-related activity of other cortical areas, we also recorded from two prefrontal regions: prearcuate gyrus (PAG) in lateral frontal cortex and supplementary eye field (SEF) in dorsomedial prefrontal cortex. The recordings from PAG were performed from monkey N with a chronically implanted 96-channel microelectrode array (electrode length=1 mm; spacing=0.4 mm; Blackrock Microsystems, Salt Lake City, UT). We performed 11 recording sessions, which yielded 84 ± 10 (mean±s.d.) simultaneously recorded units per session. The recordings from SEF were performed from monkey O with either single tungsten microelectrodes (FHC) or 16-channel linear array probes (Plexon). In 9 recording sessions, we isolated 34 SEF units. SEF was identified based on stereotactic coordinates, saccade selectivity, and evoked-saccades with low-current microstimulation (< 50*μA*).

### Behavioral data analysis

We sorted the stimulus strengths for each task into five levels. For the motion task, we used motion coherence to define the strength levels, compatible with past studies (Kiani et al. 2008; Kiani and Shadlen 2009; Shadlen and Newsome 2001): 0–3.2%, 6.4%, 12.8%, 25.6%, and 51.2% coherence. For the face task, we used the average morph level across stimulus-mask cycles in each trial: 0–5%, 5–15%, 15–25%, 25–50%, 50–100% morph. Due to the stochastic nature of the stimuli, this average morph level could differ from the nominal morph level. Specifically, in addition to changing the stimulus along the relevant axis for the categorization task, the fluctuations also shifted the stimulus along the irrelevant axis. Using the average morph level enabled us to quantify the veridical stimulus strength along both axes and explore their effects on the behavior and neural responses. Apart from this benefit, we confirmed that defining the strength levels based on the nominal morph level did not critically change any of our results. Similarly, we confirmed that defining the stimulus strength levels of the motion task based on average motion energy — a measure of veridical stimulus strength (Adelson and Bergen 1985; Kiani et al. 2014a) — did not change any of our conclusions. The same stimulus strength levels were used for both behavioral and neural data analyses.

Note that the stimulus strength could be in favor of one choice or another. Therefore, for analyses that depended on the exact choice made by the monkey (e.g., T_in_ vs. T_out_), we expanded the definition of stimulus strength levels to add a sign to the numbers above, where positive and negative strengths indicate stimuli supporting the T_in_ and T_out_ choices, respectively. Throughout the paper, we use consistent colors to show different stimulus strength levels in the figures, with solid lines or filled symbols depicting positive strengths and dashed lines or hollow symbols depicting negative strengths.

Psychometric functions (Fig. 1C, E) were calculated as the probability of making correct choices — choices compatible with the sign of the stimulus strength level — as a function of stimulus strengths. Psychophysical thresholds were defined as the stimulus strength that yielded 81.6% correct choices. Thresholds were estimated by fitting a logistic function to the choice data (similar results were obtained with cumulative Weibull fits). To assess the effect of stimulus duration on behavioral performance, we divided trials into 6 stimulus duration quantiles, calculating the psychophysical threshold separately for each quantile (Fig. 1D, F).

To test if the behavioral performance is consistent with an evidence accumulation mechanism, we fit a drift diffusion model (DDM) to the data. In the DDM, signed momentary sensory evidence accumulated over time to create the decision variable (DV). The process continued until the DV reached either an upper or a lower bound, or all the available sensory evidence was integrated. The bound that was reached first dictated the choice. But if the accumulated evidence failed to reach a bound by the end of the trial, the sign of the DV determined the choice. We fit the DDM to individual monkeys’ behavior using a maximum-likelihood procedure. Details of the method are described elsewhere (Okazawa et al. 2018). The DDM had two free parameters in the motion task: sensitivity and bound height. Sensitivity, *k*, determined the linear scaling of the mean momentary evidence in the model with signed stimulus strength, *s*. Therefore, the drift rate of the diffusion process at each moment, *μ*(*t*), was equal to *ks*(*t*). Since stimulus strength was defined based on motion coherence, *s*(*t*) was a constant in the model. The bound height, *B*, determined the amount of evidence that had to be accumulated to reach the upper (+*B*) or lower (−*B*) bound. Because the drift rate and bound height scale with the s.d. of momentary evidence in the model, we set the variance of the momentary evidence to 1 to allow unique solutions for the model fit.

For the face task, we allowed different sensitivity parameters for the three informative features in each categorization task: *μ*(*t*) = *k_e_s_e_*(*t*) + *k_m_s_m_*(*t*) + *k_n_s_n_*(*t*), where *s_e_*(*t*), *s_m_*(*t*), and *s_n_*(*t*) are signed morph levels of eyes, nose, and mouth along the relevant axis for categorization at time *t*. We did not observe any notable influence of morph levels along the irrelevant axis on the monkey’s choices, within the tested ranges in our tasks (Fig. 6B). We therefore excluded them from the equation for drift rate to limit the number of model parameters. In a reaction-time version of the task performed by human subjects, we established that sensitivity parameters were largely constant over time (Okazawa et al. 2020) and that there was minimal loss of information (leak). We do not present these model variants in this paper as they do not directly pertain to our main conclusions. However, we note that we do not find substantial evidence for non-stationary sensitivity or the presence of a significant leak in the monkey data, suggesting a similarity of the decision-making process across species.

The monkeys that performed the same task had consistent behavior and neural responses. We therefore combined data across subjects in the figures. But when appropriate, we show individual monkeys’ results too (e.g., Fig. S2). The behavioral plots in Fig. 1C-F combine data from all monkeys performing the respective tasks. The model fits (gray lines) are the average of fits for individual monkeys. Fig. 1C-D combines data and fits across both monkeys and both face categorization tasks. Fig. 6B shows the behavior separately for the species and expression categorization tasks. Since the veridical stimulus strengths varied along both task-relevant and irrelevant axes, we plotted two psychometric functions, one for each axis. Both curves in the figure are from the DDM explained above. The model fits do not include any degree of freedom for stimulus strengths along the irrelevant morph axis, as explained above. The non-significant changes of model accuracy along the irrelevant morph axis are due to random stimulus fluctuations along the task-relevant axis that do not totally average out for the limited number of trials in the dataset.

### Neural data analyses

Peri-stimulus time histograms (PSTHs) shown in Figs. 2A-B, 5A, 5C, 7A, S2, S4A, S5D, S8A and S9C, E were smoothed by convolution with a 100 ms boxcar filter. The PSTHs were aligned to the stimulus onset and cut off at 300 ms after stimulus offset to focus our analysis on the responses pertaining to the decision formation based on sensory inputs. The same convention was used in subsequent analyses. When plotting the PSTHs, we focus on correct trials only to be able to examine changes of firing rate with stimulus strength for the same choice except for Fig. S2E.

The firing rates shown in Figs. 2C-D and S8B were calculated by counting the number of spikes during decision formation (250-600 ms after stimulus onset) in correct trials. To combine data across units with different ranges of firing rates, we z-scored the firing rates of each unit by subtracting its mean firing rate and dividing by the standard deviation of firing rates across all trials. We then calculated the mean of z-scored firing rates of each unit for each stimulus strength and averaged them across units. To quantify the relationship between the firing rates, *r*(*s*), and the stimulus strength, *s*, we performed linear regressions independently for T_in_ and T_out_ choices:

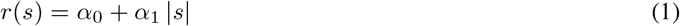

where |*s*| is the absolute (unsigned) stimulus strength. The regression coefficients (*α*_0_ and *α*_1_) were determined for each unit. The lines in Figs. 2C-D and S8B were generated from average *α*_0_ and *α*_1_ across units. The significance of slopes across the neural population was determined based on a two-sided *t*-test on the distribution of single unit slopes.

To examine if the difference of T_in_ and T_out_ firing rates increased with stimulus strength, we used the following linear regression:

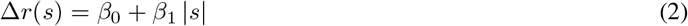

where Δ*r*(*s*) is the differential firing rate between correct T_in_ and T_out_ choices for the unsigned stimulus strength |*s*|. Fig. 2F shows the distribution of the slope coefficient (*β*_1_) across the recorded units.

We implemented a neurometric analysis to quantify the accuracy of an ideal observer that uses the responses of LIP neurons to discriminate the stimuli with similar strength on the two sides of the category boundary in each task (Britten et al. 1992; Shadlen and Newsome 2001). The ideal observer accuracy equals the area under the Receiver Operating Characteristic (ROC) curve of the distribution of spike counts for the two competing stimuli. Figure S2D shows the results for different stimulus strengths using a 100ms sliding window. To remain conservative, we did not conditionalize spike count distributions on correct choices in this analysis. Because conditioning on correct choices would lead to higher areas under ROC for weaker stimuli, it would further reduce the dependence of the ideal observer’s accuracy on stimulus strength, bolstering our conclusion that the difference of the activity of pools of neurons selective for the two choices fails to fully explain the representation of the DV in the face task.

The PSTHs for the reaction-time version of the motion task (Fig. 7A) were calculated from 100 ms before stimulus onset to the median reaction time for each stimulus strength. A 50 ms window before saccade onset was excluded from each trial so that the PSTHs were not influenced by the motor burst of the neurons. Using a more conservative exclusion window (100 ms before saccade onset) did not critically change the results. For both the PAG and SEF units, target selectivity was determined using neural responses during a presaccadic period (–300 to 0 ms from saccade onset) and the choice target that evoked higher activity was defined as the putative T_in_. Responses in memory- or visually-guided saccade tasks corroborated our designations.

In the following sections, we explain analyses for characterization of population response patterns. We use these population analyses as a vehicle to drive intuition and develop insights, noting that because these methods build on single cell responses, their conclusions can also be derived from careful analysis of single cells across the population.

### Principal component analysis of neural data

We performed principal component analysis (PCA) to visualize the neural population response manifolds (Figs. 2G-J, 5A-D, 6C, 7B, S4, S5, S8C and S9D, F). PCA was performed across units using trial-averaged PSTHs. Each unit contributed multiple PSTHs, one for each combination of choices and stimulus strength levels. The PSTHs focused on the period of decision formation and spanned 250-600 ms after stimulus onset. We detrended the PSTHs of each unit by subtracting the average unit PSTH across all stimulus strengths and choices to focus on modulations of activity associated with task parameters. We confirmed that the detrending did not critically change the observed geometry of neural manifolds. We then concatenated the detrended PSTHs of each unit into a response vector for that unit and combined the vectors of all units into a population response matrix with *T* × *C* rows and *N* columns, where *T* is the number of time points in each PSTH (1 ms resolution), *C* is the number of conditions (5 stimulus strength levels × 2 choices, T_in_ and T_out_), and *N* is the number of units. Only correct T_in_ and T_out_ choices were used for these analyses because error responses were rare for high stimulus strengths. A tiny fraction of elements in the population response matrix were missing (< 0.1%) due to low numbers of trials for some units. The missing data were replaced with the average response of the corresponding unit. The first three principal components (PCs) explained 78% of the total variance of the full *N* dimensional neural space in the motion task and 54% of the total variance in the face task. The analysis included all units recorded either simultaneously or separately. We confirmed that similar results were obtained using simultaneously recorded units only (Fig. S5C).

Because we have aligned the PSTHs of LIP neurons with respect to their RFs (Fig. S3A; PSTHs were defined with respect to T_in_ and T_out_, not the right-left location of targets), the PCA effectively captures the structure of population responses within a group of neurons having a shared choice preference. Therefore, the observed manifold geometry arises from the variability of neural responses within that group of neurons (see Fig. S3B for intuition). The finding, however, also extends to a neural population with diverse choice preferences. We performed PCA for a pseudo-population created by inverting the preferred choice of a random half of the recorded units. The resulting manifold geometry lead to conclusions similar to our main results (Fig. S3C).

The neural population response for each stimulus strength level at each time can be depicted as a point in the three-dimensional (3D) PC space (Figs. 2G-J, 5A-D, 6C, 7B, S4B-D, S5, S8C and S9D, F). The data points in the figures were calculated based on spike counts within a 100 ms window centered at the specified times in each figure. To draw a manifold that captures the population responses for different stimulus strength levels (gray lines), we fit the projection of population responses along each PC axis with a cubic smoothing spline as a function of stimulus strength (*csaps* function in Matlab). The s.e.m. of the population response projections along each PC axis (error bars in Figs. 2G-H, 7B, and S8C) was computed using a bootstrap procedure. We re-computed the mean firing rate of each unit for each stimulus strength level and time by randomly sampling trials with replacement. Then, we projected the resulting population response patterns onto the 3D PC space. This procedure was repeated 1,000 times, and the standard deviations of the projections were calculated and used as the s.e.m. of the data points in the plots.

To rule out the possibility that the population response manifold is actually a straight line that looks curved due to noise in the measured neural responses, we performed three tests. The first two examined the consistency of the direction of the curvature using a cross-validation procedure, and the third tested whether the observed extent of the curvature could arise solely from neural noise. For the analyses in the paper, the three tests always provided consistent results.

Stochastic variability of neural activity could indeed cause changes in the curvature of the population response manifold, but these noisy changes are in random directions in the state space and would lack consistency. In our cross-validation tests, we divided the trials of individual units into random halves and derived a manifold for each half. In the first test, we directly compared the directions of the major axes of the two manifolds, 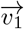 and 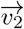. The major axis of each manifold was defined as the vector between the midpoint of the two ends of the manifold and the apex of the manifold. The angles between the two major axes were quite small, 11.9° ± 6.0° in the motion task and 9.3° ± 4.9° in the face task (mean ± s.e.m., calculated over 1,000 iterations of random splitting of data). These angles were significantly smaller than expected from noise (90°), indicating strong consistency in the direction of manifold curvature.

In our second cross-validation procedure, we adopted a more robust approach that averaged the direction of curvature of all locations of the manifold for the second half of the data with respect to the major axis of the manifold from the first half. For the 1D manifold 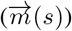 in the state space at the location corresponding to stimulus *s*, its curvature is:

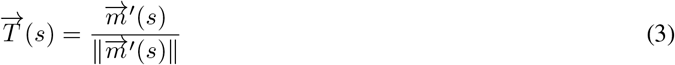

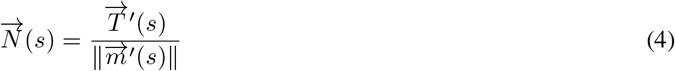

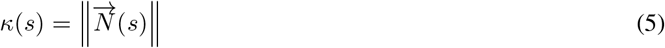

where 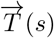 is the unit tangent vector, 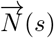 is a vector normal to the curve, and the length of 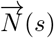 equals the curvature, *κ*(*s*), at the location corresponding to stimulus *s*. ||.|| is the *L*2 norm. The direction of 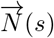 determines the direction of curvature in the state space. The dot product of 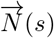 with 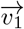 determines how well their directions match each other:

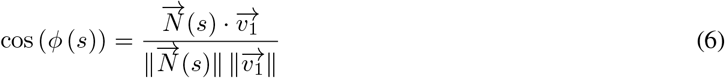

where *ϕ*(*s*) is the angle between 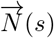 and 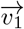 at *s*. cos (*ϕ* (*s*)) was computed at every point on the manifold and then averaged over the full extent of the manifold. Positive values of 〈cos (*ϕ* (*s*))〉_*s*_ indicate that the direction of curved manifold for the second half of data is consistent with the first half. We tested if this value is significantly greater than zero using a bootstrap procedure (iterations, 1,000).

For our third test of the significance of manifold curvature, we compared the curvature of the actual neural data with that of simulated data from a linearized response manifold with similar conditional response variability to the actual data (Fig. S5A). The simulated data were generated by straightening the underlying manifold in the actual data. We first redefined the mean firing rates of each unit for different stimulus strengths by linearly interpolating between firing rates for the strongest stimuli supporting T_in_ and T_out_ choices. This interpolation ensured the linearity of the simulated response manifold. For each stimulus strength, we defined the distribution of simulated responses of a unit as a mean-adjusted version of the true distribution of unit responses, such that variance and shape of stimulus-conditioned response distributions remained unchanged. We simulated individual trials by randomly sampling from these adjusted distributions for the same number of trials that each unit was recorded for. Repeating this process for all the units generated a simulated dataset with the same number of units, trials, and conditional response variability as the actual data. We created 1,000 simulated datasets and derived the manifold for each of them using the procedure explained above. Then we calculated the curvature at each point of each manifold using Eq. 5 and quantified the overall magnitude of the curvature as the average curvature across the manifold. Finally, the magnitude of the manifold curvature in the data was compared with the distribution of magnitudes obtained from the simulated data.

In the PC space, the projected population average firing rate changed along a linear axis (straight line in Figs. 2I-J, 6E, and S4B), since PCA is a linear dimensionality reduction procedure. A vector that corresponds to this linear projection axis can be readily derived from the PC coefficients:

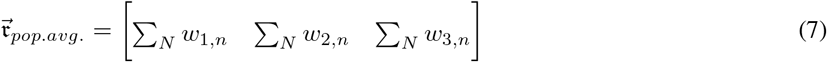

where *w_i,n_* is the coefficient of neuron *n* for the *i*-th PC. In Fig. 2I-J, to illustrate the population average firing rates corresponding to locations on the manifold, we projected a perpendicular line from each point on the manifold to the axis of the average firing rate in the 3D PC space. Due to limitations of 2D illustrations, however, these perpendicular projections in the 3D space appear to have angles different from 90° in the figure.

For the reaction time tasks, PCA was performed as described above but using the detrended PSTHs from 250 ms to 550 ms after stimulus onset. The PAG results in Fig. 7 were generated using the simultaneously recorded units in one session (*n* = 99). Similar results were obtained in all recorded sessions.

### Locating the DV and stimulus difficulty axes in the population response state space

The curved population response manifold suggests the presence of linear axes in the state space that encode the decision variable (DV) and stimulus difficulty (Fig. 3A). To identify the best encoding axes, we performed an orthogonal canonical correlation analysis (CCA) (Cunningham and Ghahramani 2015) (Fig. 3B). CCA finds linear transformations that maximize the correlation of two matrices with each other. One matrix in our analysis was the neural population response, **R**, consisting of the trial-averaged, detrended PSTHs of the recorded units. The population response matrix was constructed as we explained for the PCA analysis but was limited to a shorter window (350–450 ms after stimulus onset) in order to allow approximation of the DV and stimulus difficulty based on task parameters. Neural activity around 400 ms after stimulus onset well reflects the decision formation, but similar results could also be obtained at earlier or later times (e.g., 300 ms, 500 ms, or 600 ms). The second matrix in our analysis was the task parameter matrix, **P**, which approximated the DV and stimulus difficulty associated with each PSTH as *s* (signed stimulus strength) and − |*s*|, respectively. For simplicity, we assumed a constant DV and difficulty during the short PSTH snippet included in the response matrix. The average DV across trials of a particular stimulus strength is proportional to the signed stimulus strength in our tasks, especially at early periods of the decision-making process when the decision bound is not yet reached for the majority of trials because the behavior could be successfully modeled with linear integration of stimulus strength (Figs. 1C-F, S1). The proportionality constant, however, changes with time, as the DVs for different stimulus strengths diverge from each other (Beck et al. 2008; Kiani et al. 2014b; Peixoto et al. 2018). It is therefore important to limit the duration of analysis window to ensure reliable CCA results, while keeping it long enough to allow accurate estimation of mean firing rates in each condition. A 100 ms window met the analysis requirements, but our conclusions were not critically dependent on it. Further, the exact time of the analysis window had minimal impact on our results as long as it reasonably overlapped with decision formation (we tested 350–600 ms after stimulus onset). More complex methods in which we inferred the DV and task difficulty using the exact stimulus fluctuations corresponding to the analysis window and the DDM yielded similar results.

The CCA seeks projection vectors, 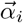 and 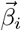, that maximize the correlation of canonical variables 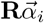 and 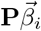. After finding the first pair of canonical variables, the analysis finds a second pair of canonical variables uncorrelated with the first. Orthogonal CCA also ensures that the projection vectors are orthogonal (Cunningham and Ghahramani 2015). Combining the projection vectors yield 2D transformation matrices 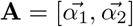 and 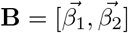, which we used to project the neural responses to the 2D subspace defined by the linear DV and difficulty axes:

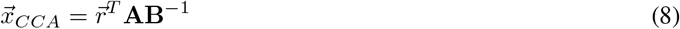

where 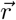 is the population response vector and 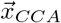 is the position in the 2D subspace. In Figs. 3, 4, 6H, and S6, 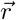 consisted of firing rates of the neurons in 100 ms windows. For the CCA results presented in the paper, we reduced the dimensions of **R** to 10 using PCA to attempt denoising the population responses prior to the CCA. However, the denoising or the number of PC dimensions used for denoising was not critical for our conclusions (we tested a wide range of dimensions from 10 to 40; 10 dimensions explained 89% and 76% of total variance in the motion and face tasks, respectively).

All figures and analyses that depended on CCA were cross-validated. The cross validation was implemented by using a random half of trials as a training set to compute the transformation matrices, and then applying them to the other half of trials (test set). In Figs. 3B-D and 4C-E, the s.e.m. of the data points were estimated using a bootstrap procedure within the test trials (iterations, 1,000). The gray curves in Figs. 3B and 4C, which estimated the manifold spanning different stimulus strengths, were computed by fitting the projection of neural responses along each dimension using a second-order polynomial function of stimulus strength.

To plot the time course of neural responses along the two axes defined by CCA (Fig. 3C, D), we further split the test trials based on the monkey’s choice (correct or error) or based on stimulus difficulty (easy or difficult, specified as stimulus strengths greater or less than 20%). When computing the choice-dependent neural responses (Fig. 3C), all stimulus strengths were combined except for 0%, for which the correct choice was undefined. When computing the difficulty-dependent neural responses (Fig. 3D), we used only correct trials, thus keeping the monkey’s choice identical between the easy and difficult conditions.

To examine the relationship between the neural responses and the monkey’s confidence, we analyzed a subset of neurons (70/129) recorded during the motion task with post-decision wagering (see the section on Behavioral tasks). Bailing out of the motion direction discrimination by choosing the sure-bet option indicated low confidence. We first performed CCA on a random half of trials with correct choices (training set) to find the best encoding axes for the DV and stimulus difficulty. Then, we projected neural responses of the remaining trials on these axes, independently for correct, error, and T_s_ choices. In Fig. 4C, we adjusted the sign of the DV axis projections such that positive and negative values indicated DVs in favor of the correct and error choices, respectively. To quantitatively test the relationship between the neural responses and the monkey’s choices (Fig. 4D, E), we computed the residual activity associated with each choice (correct, error, and T_s_) by subtracting the mean projection of all test trials with the same stimulus strengths along each axis. We then averaged the residual responses across different stimulus strengths to achieve better visualization and boost analysis power. For these analyses, we used only trials with low stimulus strengths (3.2%–12.8%), where error and T_s_ choices were present in all sessions. Because correct and error choices were undefined for 0% coherence trials, they were not included.

### Testing task dependency of neural responses

To examine how task contexts influenced LIP activity, we compared the neural responses during the species and expression categorization of the same face stimuli (Fig. 6). The analysis used a subset of units (n = 28) recorded in both contexts in the same sessions. To create the PC space common to both categorization tasks, we used stimuli present in both tasks (purple region in Fig. 6A). The stimuli were grouped into four strength levels (−15%–−5%, −5%–0%, 0%–+5%, +5%–+15%) for each stimulus axis (species and expression). This yielded 32 conditions (4 species levels × 4 expression levels × 2 task contexts). We computed PSTHs of correct trials in these conditions and performed PCA using the same procedure described earlier.

In Fig. 6D, we plotted the PC scores of each stimulus strength (4 × 4 levels) in a window 350-450 ms after stimulus onset. The significance of differences in PC scores between the tasks was determined using a permutation test. For each stimulus level, we shuffled the trial labels between the two tasks (preserving the number of trials) and computed PC scores of the two shuffled groups. We then evaluated the similarity of the two groups of PC scores by calculating the Pearson correlation coefficient across stimuli and PC dimensions. We repeated this procedure 1,000 times to establish the null distribution and compared it against the observed correlation coefficient to report the *p*-value.

In Fig. 6C, we selected the top three PCs derived from the above analysis and projected neural responses calculated for our standard stimulus strength levels in each task as explained in earlier sections. Here, the statistical significance of the difference of the manifolds between the two tasks was evaluated using a likelihood ratio test. We fit the PC scores of each dimension using a second-order polynomial function of stimulus strength, either simultaneously, using the same coefficients for both tasks, or separately, using different coefficients for each task. The goodness of fits was quantified as the likelihood of data points given the manifolds under a Gaussian noise model. The noise covariance matrix was estimated using a bootstrap procedure (iterations, 1,000). The likelihood ratio test determined whether the likelihood of separate manifolds for the two tasks was significantly larger, taking into account the difference in the degree of freedom between the two fits.

The task dependency of neural responses influenced how the DV was encoded by the population. To examine the extent of this effect, we first determined the DV axis in each task context using the CCA procedure described earlier, and then we computed the angle between the DV axes of the two face categorization tasks in the 3D PC space (*θ* in Fig. 6E). We used the neural responses in a window 350-450 ms after stimulus onset, but the results were largely invariant to the exact choice of analysis window during decision formation. *θ* was defined with respect to the axis of average firing rate (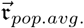 in Eq. 7) such that positive values indicated a greater angle between 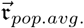 and the DV axis in expression task, compared to that in species task. Hence, when the DV axes in the two categorization tasks differ, *θ* should be either significantly greater or less than zero. We used a bootstrap procedure to test the statistical significance of *θ* (iterations, 1,000).

We quantified the extent to which the angle between the DV axes (*θ*) affected the linear readout of the encoded DVs. We projected the neural responses recorded in one task context on the DV axis derived for the same or different context (Fig. 6H; the results of the expression task are plotted). The projected responses were fit with a logistic curve as a function of stimulus strength (*s*):

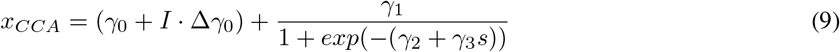

where *γ_i_* are regression coefficients and *I* is an indicator variable (0 for using the DV axis from the same context and 1 for using the DV axis from the different context). Δ*γ*_0_ captures the change of bias caused by not adjusting the DV axis based on task context. The significance of Δ*γ*_0_ was examined using a likelihood ratio test that compared the likelihood of models with and without Δ*γ*_0_. The likelihoods were estimated using a bootstrap procedure (iterations, 1,000).

The analyses above used only the neurons recorded in both task contexts. Because the rest of units (n = 104) were recorded for only one task context, their DV axes could not be directly compared. However, the axis of average firing rate 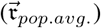 could be used as a common reference. We reasoned that, if a brain area encodes the DV in a context-independent manner, a population of neurons recorded from this area would have a fixed angle between the DV axis and the axis of average firing rate (i.e., *ω_spc_* = *ω_exp_* in Fig. 6E). We therefore tested if the angle between the DV axis and 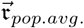 differed between the populations recorded for each face categorization task (Fig. 6G). The s.e.m. of the angles and statistical significance of the angle differences were calculated using a bootstrap procedure (iterations, 1,000). Repeating these tests in higher dimensional state spaces provided similar conclusions.

## Acknowledgements

The authors would like to thank Long Sha, Michael Waskom, Saleh Esteki, Bianca Sieveritz, Mike Shadlen, Stefano Fusi, and Alex Pouget for helpful discussions and comments on earlier versions of the manuscript. This work was supported by the Simons Collaboration on the Global Brain (grants 542997 and 543009), McKnight Scholar Award, Pew Scholarship in the Biomedical Sciences, and National Institute of Mental Health (R01 MH109180-01). G.O. was supported by post-doctoral fellowships from the Charles H. Revson Foundation and the Japan Society for the Promotion of Science. C.E.H. was supported by National Eye Institute Training Grant in Visual Neuroscience (T32 EY007136).

## Supplementary figures

**Figure S1:**
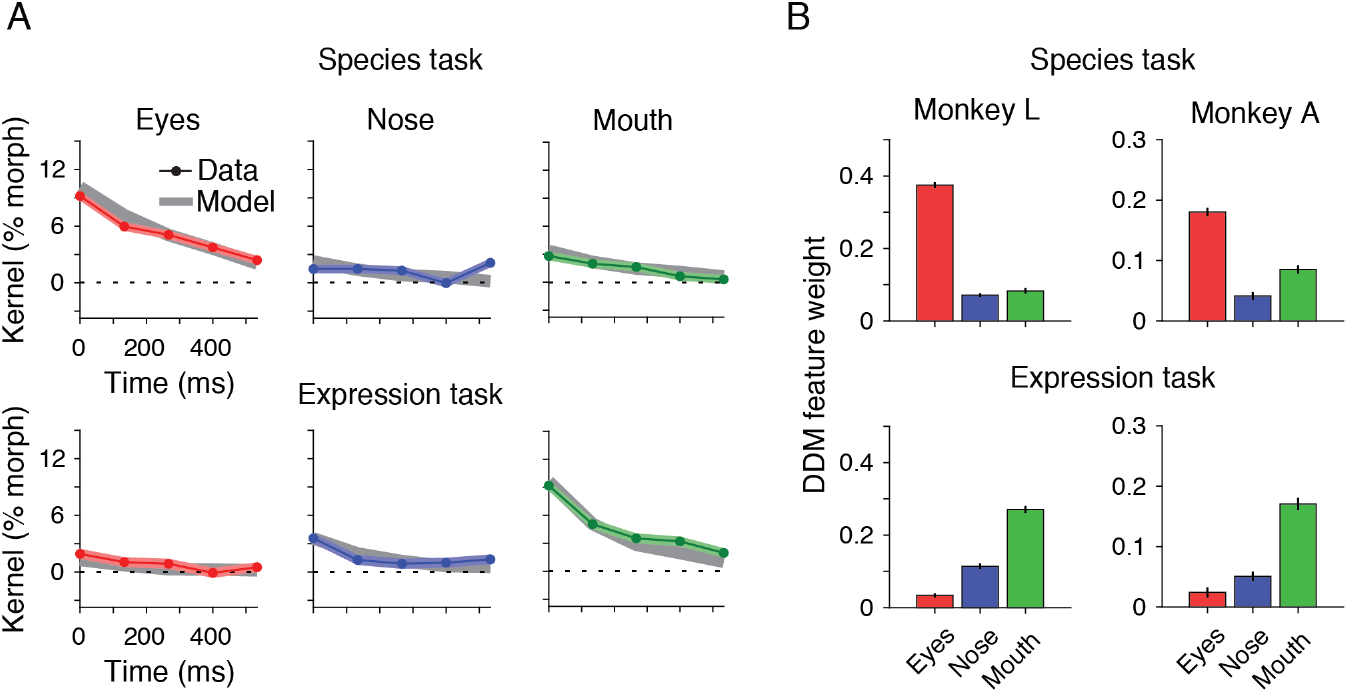
Psychophysical kernels of the face task support bounded evidence accumulation with task-dependent weighting of facial features. (**A**) In the face task, morph levels of the three informative facial features (eyes, nose, mouth) randomly fluctuated every 106.7 ms around a mean morph level on each trial (nominal morph level). These fluctuations allowed us to perform psychophysical reverse correlation (Ahumada 1996) to examine spatiotemporal weighting of features for decision making. Psychophysical kernel, *K_f_* (*t*), for each facial feature *f* at time *t* was calculated as the difference of average fluctuations of morph levels conditional on the monkey’s choices:

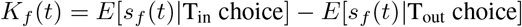

where *s_f_* (*t*) is the morph level of feature *f* at time *t*. We used trials with nominal morph levels less than 20%, where stimulus fluctuations were potent enough to switch the monkey’s decision. To calculate a kernel across different morph levels, we subtracted the nominal morph level of each trial from the actual morph levels presented on the trial and then averaged these residuals across trials. Colored lines show the resulting kernels for species categorization (top row) and expression categorization (bottom row). Kernel amplitudes were different across features and tasks, indicating that (1) the three informative features differentially contributed to the monkey’s decisions in each task, and (2) the feature weights varied across tasks. Species categorization relied most heavily on the eyes but expression categorization on the mouth. Further, the kernels decreased over time in our task design, compatible with bounded integration of evidence, although such declines could also arise from other mechanisms (Okazawa et al. 2018). To quantify the weights of the three features for the monkey’s choices, we approximated the decision-making process with a drift diffusion model (DDM) that accommodated differential weights for the informative features along the relevant morph axis for each categorization task (see Methods). Fitting the model to the distribution of choices across trials quantitatively explained monkeys’ psychometric functions and changes of threshold with stimulus viewing duration (Fig. 1C,D). Further, it generated predictions for the shape of psychophysical kernels that closely matched the data (gray lines). The predicted kernels were calculated by creating new stimulus sequences that were not used for fitting the model and simulating the DDM for these stimuli with the fitted model parameters. The predicted kernels accurately explained the data (species categorization, *R*^2^ = 0.97; expression categorization, *R*^2^ = 0.98). (**B**) Feature weights of the DDM fits in the two face categorization tasks. Error bars are s.e.m.

**Figure S2:**
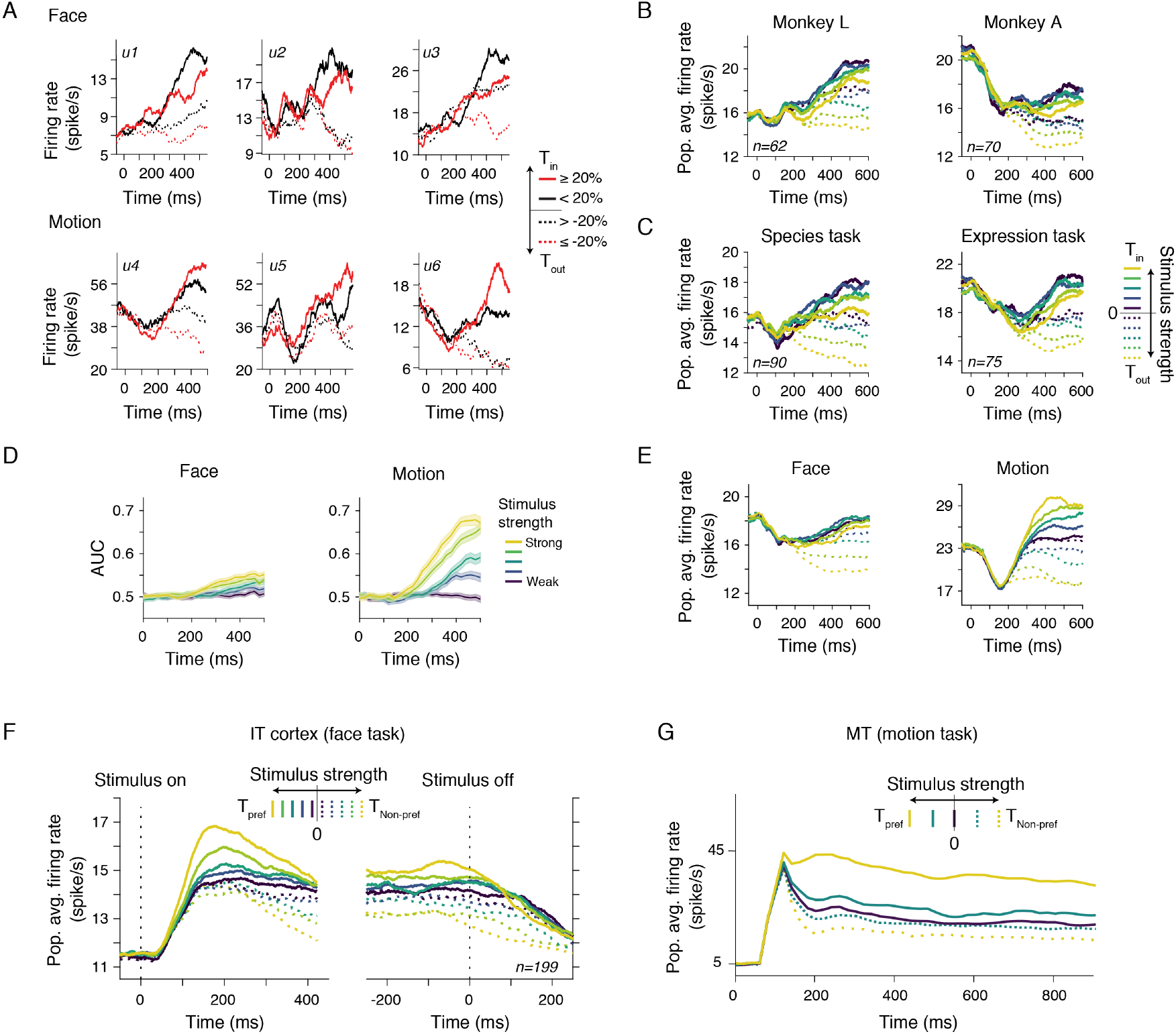
Reversed ordering of T_in_ PSTHs with stimulus strength was consistently present across units, monkeys, and different categorizations in the face task. (**A**) Units recorded in the face task (top row) showed a reversed order of T_in_ PSTHs for different stimulus strengths compared to those recorded in the motion task (bottom row). We show three example units from each task. For clarity, trials were divided into two difficulty levels (≥ 20%, red, and < 20%, black) for T_in_ (solid) and T_out_ (dashed) choices. The PSTHs show only correct responses and are smoothed with a 100 ms boxcar filter. (**B**) Population average PSTHs for individual monkeys. Conventions are the same as in Fig. 2A. (**C**) Population average PSTHs for species and expression categorizations in the face task. (**D**) Ideal observer accuracy for discriminating stimuli with equal strength on the two sides of category boundary based on individual unit responses in the face and motion tasks. The ideal observer accuracy for each time and stimulus strength was quantified as the area under the Receiver Operating Characteristic (ROC) curve for the distribution of spike counts in a 100 ms window (AUC: Area Under Curve). The lines in each panel show the average AUC across the recorded units. The shading indicates s.e.m. For low stimulus strengths, the spike count distributions are largely overlapping and indistinguishable (AUC≈0.5). The spike count distributions move systematically apart from each other for stronger stimuli in the motion task, leading to ideal observer accuracies much larger than chance (AUC≫0.5), especially for longer viewing durations. These results match the systematic improvement of the monkey’s performance with stimulus strength and duration (Fig. 1E-F) and are expected from the monotonic ordering of PSTHs as a function of stimulus strength (Figs. 2B and S2E) in the motion task. In contrast, stronger stimuli are associated with a much smaller increase in the ideal observer’s accuracy in the face task, as expected from the non-monotonic ordering of the PSTHs with stimulus strength (Figs. 2A and S2B,C,E), and are incompatible with the monkey’s performance in the task (Fig. 1C-D). Since the ideal observer analysis relies on the difference of responses of a pair of neurons with opposite choice preference—neuron and anti-neuron (Britten et al. 1992; Shadlen and Newsome 2001), the poor separation of AUC lines with different stimulus strengths in the face task suggests that the decision variable could not be encoded as a simple difference of responses of two pools of action selective neurons. Past studies in LIP conditioned the ideal observer analysis on choice (e.g., Shadlen and Newsome 2001). Doing so would further reduce the separation of AUC lines, bolstering our conclusion. (**E**) Population average PSTHs including both correct and error trials. Including error trials leads to different proportions of T_in_ and T_out_ choices for different stimulus strengths. A mixture of T_in_ and T_out_ choices for lower stimulus strengths moves their PSTHs toward the center of the firing rate range. Nonetheless, the reversal of the order of the PSTHs was still present in the face task. (**F-G**) Neurons in the IT cortex show monotonic tuning for stimulus strength during the face categorization tasks, similar to the monotonic tuning of MT cells for motion strength. Firing rates of MT neurons are known to monotonically change as a function of motion coherence of the random dots stimuli (G; reproduced from Britten et al. 1993). If sensory responses to our face stimuli show non-monotonic, complex tuning, integrating these responses might result in non-monotonic firing rates as observed in LIP. To exclude this possibility, we analyzed the activity of neurons in the IT cortex recorded simultaneously with LIP neurons during the face tasks. The recordings were made from face selective neural clusters in anterior IT (Tsao et al. 2003). Panel F shows the population average firing rates of 199 face-selective neurons. Stimulus strengths were sorted according to individual neurons’ preferred face category. The firing rates were monotonically modulated by the stimulus strength, providing evidence against the possibility that the reversal of firing rates in LIP is inherited from the activity of sensory neurons.

**Figure S3:**
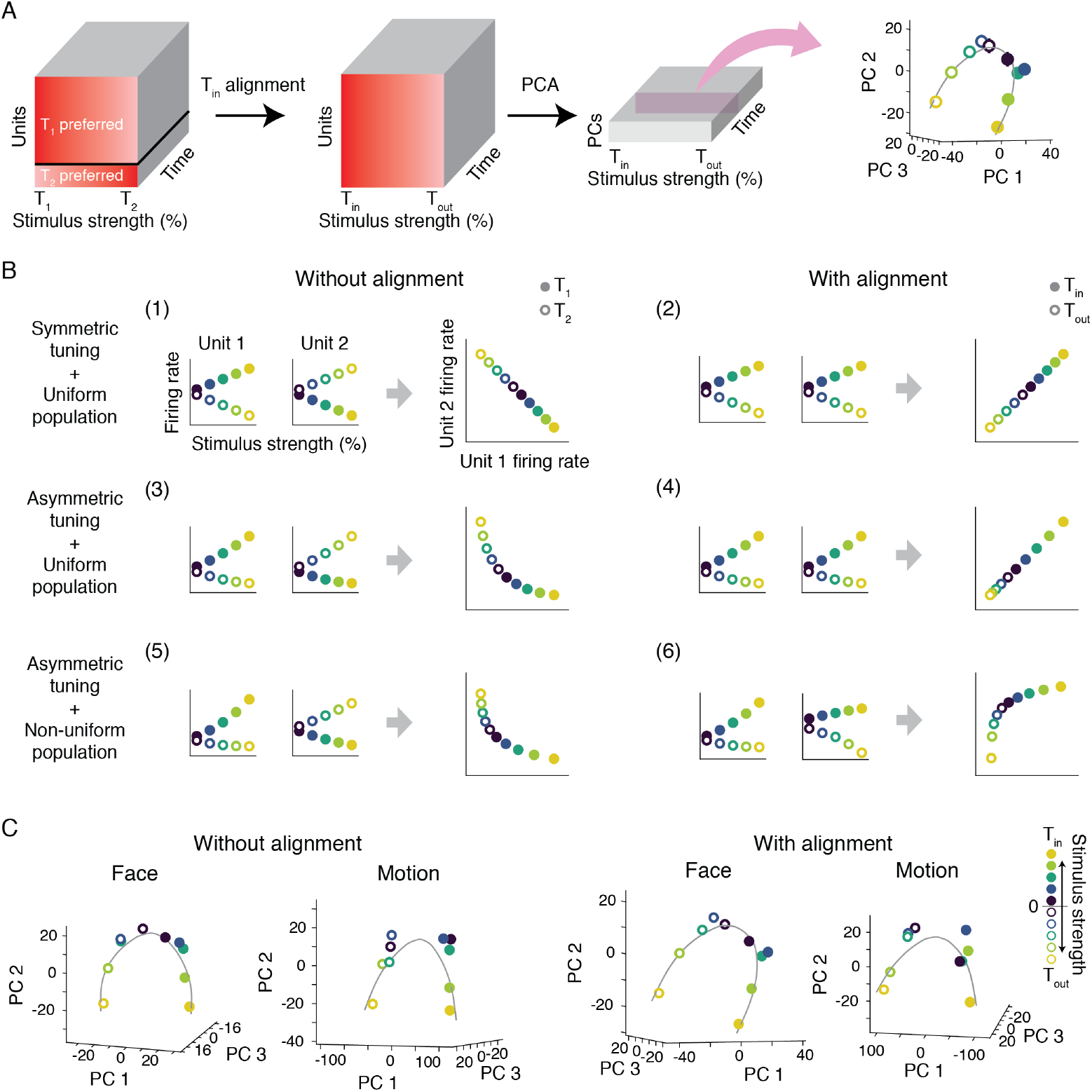
A curved manifold in the PC space arises from the variation of tuning for stimulus strength across the recorded population. (**A**) A schematic description of our principal component analysis (PCA). Our dataset included neurons preferring either of the two targets. T_1_ and T_2_ indicate the targets on the right and left side of the screen, one of which in the RF. T_in_ and T_out_ indicate the targets inside and outside the neuron RF. Signed stimulus strength in the experiment was originally defined based on the location of the targets (T_1_ vs. T_2_). For each neuron, we redefined the stimulus strength such that negative values matched T_out_ and positive values matched T_in_. Doing so enabled us to analyze the recorded units as a population of neurons with similar target preferences (i.e., aligned T_in_) (middle). In our main analyses, PCA was applied to the PSTHs after this alignment (right) to focus on the diversity of stimulus tuning across the population (see panel B). (**B**) When the preferences of neurons are aligned, a curved manifold arises from the variation of neural tuning across units. To demonstrate this point, we simulated six hypothetical scenarios for a neuron pair with opposite choice preferences (one preferring T_1_ and the other T_2_), plotting their joint manifolds before and after the T_in_ alignment. If the neurons have symmetric tuning for positive and negative stimulus strengths (slopes with similar magnitudes but opposite signs) and the tuning slopes are identical across the population, the manifold would be a straight line regardless of T_in_ alignment (B1-B2). If the single-unit tuning is asymmetric but similar across the population, the manifold would be curved before alignment (B3) but straight afterward (B4). However, T_in_ alignment does not remove the curvature caused by diverse tuning across neurons (B5-B6). (**C**) The T_in_ alignment focused our analyses on the diversity of responses across the population. However, all of our main conclusions replicate in the absence of T_in_ alignment too. The left panels show the population response manifolds without T_in_ alignment for a balanced population of neurons preferring T_1_ and T_2_. The right panels show the manifolds with T_in_ alignment.

**Figure S4:**
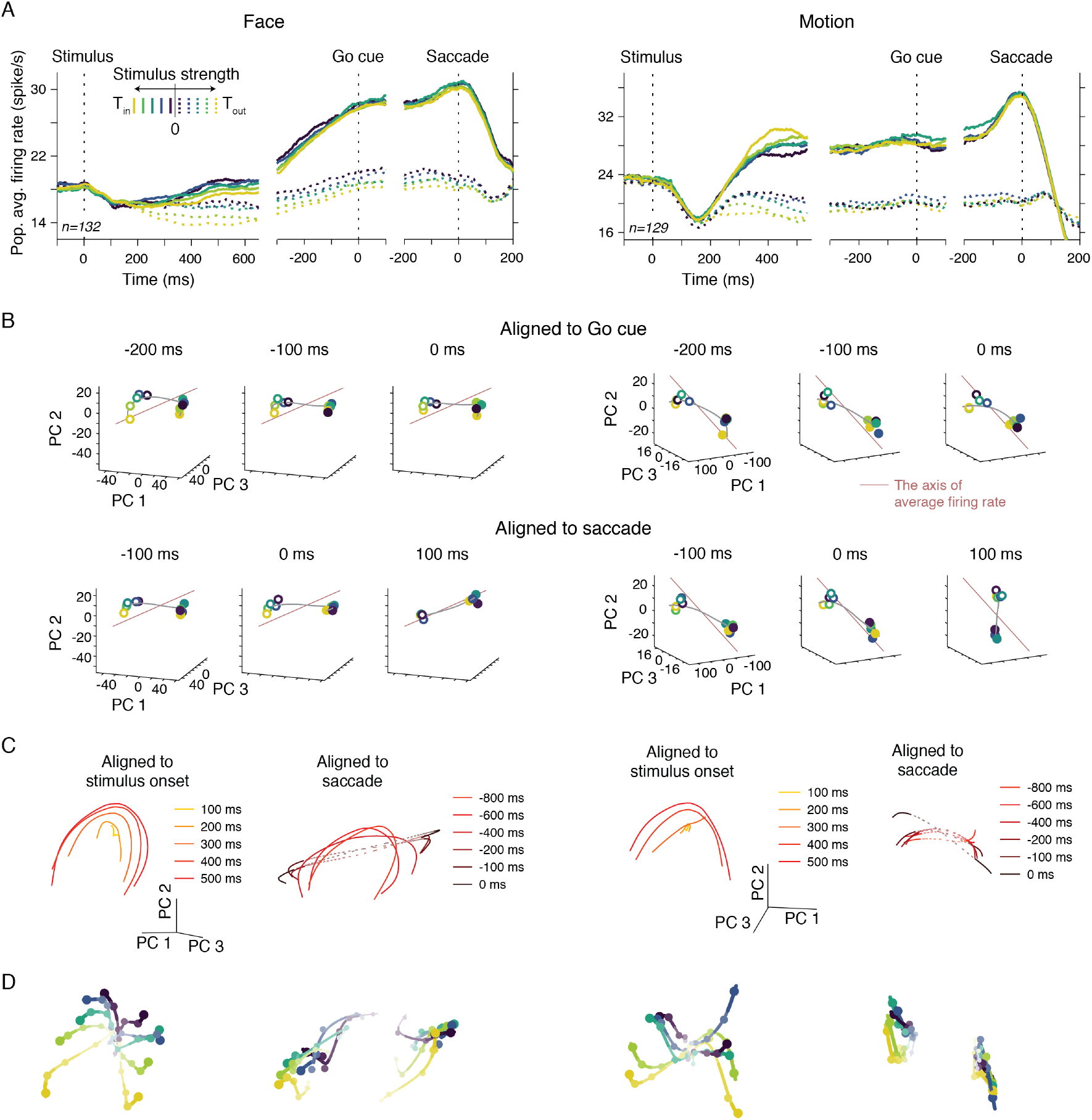
Population neural responses throughout task epochs. (**A**) Population average PSTHs aligned to multiple task events. Those aligned to the stimulus onset are the same as Fig. 2A-B. In the face task, the separation of the population average T_in_ and T_out_ PSTHs were smaller, and firing rate dynamics continued into the delay period after stimulus offset. However, in both tasks, T_in_ PSTHs converged to the same level before the saccade for all stimulus strengths, consistent with past studies (Roitman and Shadlen 2002; Shadlen and Newsome 2001). (**B**) Around the time of the Go cue and saccade, the population neural responses in the PC state space converged to one of the two points that represented the monkey’s choice. The PC space is identical to those in Fig. 2G, H. Each panel shows the responses at a particular moment aligned to the Go cue (top) or saccade onset (bottom). (**C**) Population response manifolds expanded over time after stimulus onset, remaining largely aligned during decision formation but misaligned with the action encoding axis. After the stimulus offset, population response patterns for different stimulus strengths converge onto one of the two action-encoding points in the state space, compatible with the transformation of the DV encoding to choice encoding. Regions not occupied by neural responses on the saccade-aligned manifolds are shown as dashed lines and plotted only to convey misalignment with the manifolds at earlier times. (**D**) The trajectory of population neural responses for each stimulus strength in the same PC space, aligned to the stimulus onset and saccade onset for each task. Data points represent response patterns for different stimulus strengths at different times. Lines connect data points of the same stimulus strength over time. Later times are shown with larger points and more saturated colors.

**Figure S5:**
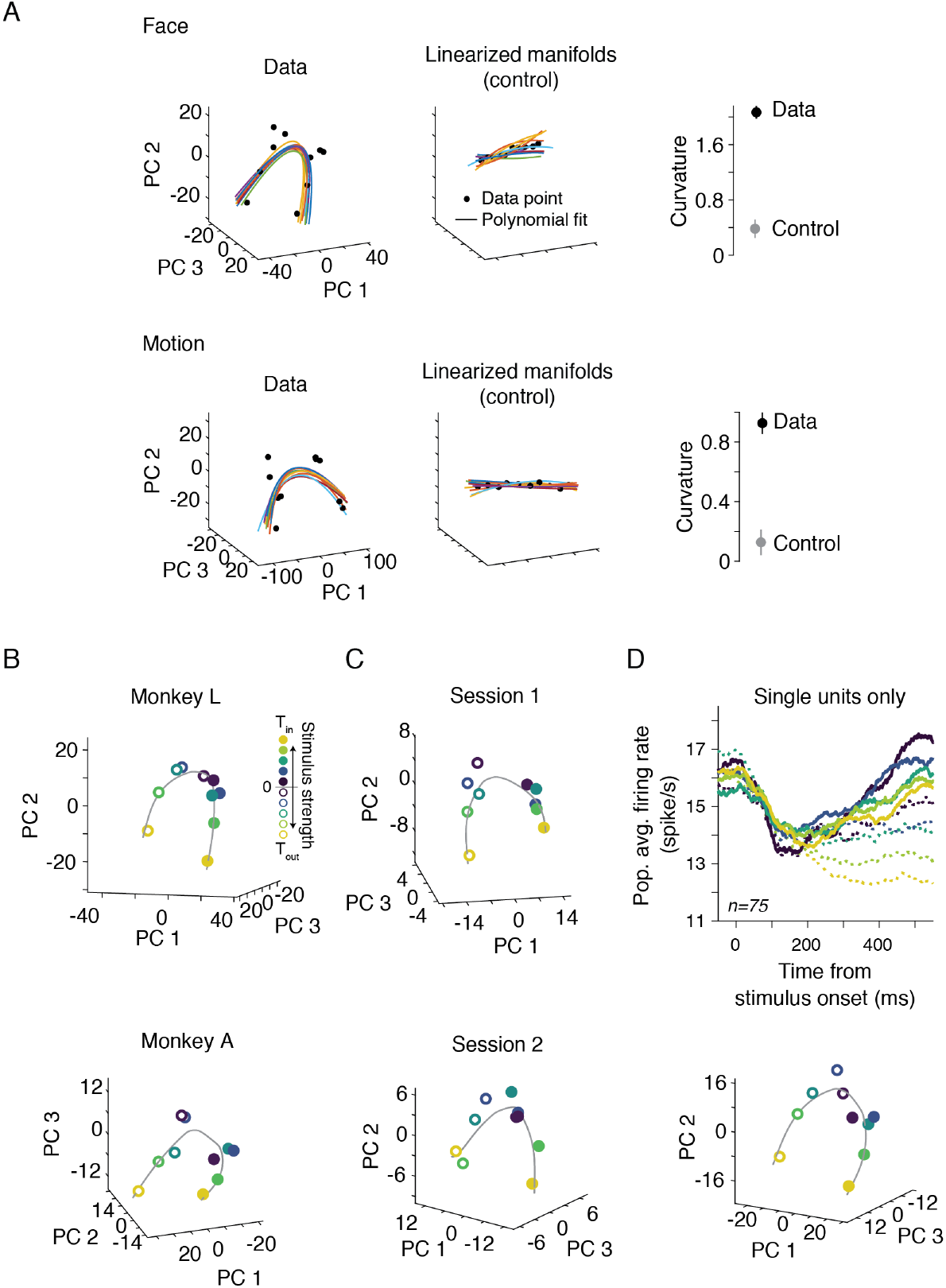
The extent of the manifold curvature in LIP population responses exceeded the curvature expected purely from neural response variability. (**A**) To confirm the significance of the manifold curvature, we compared the magnitude of curvature in the actual neural responses with that of simulated data that had the same response variability as the actual data but were constrained to a linearized manifold (see Methods). The linearized manifolds were generated by adjusting mean responses for each stimulus strength to be on an interpolating line between the responses to the strongest stimuli supporting the T_in_ and T_out_ choices. Population responses were then generated using these adjusted means while preserving the response variability across trials at the levels observed in the data. Response variabilities could bend the simulated manifolds due to finite sampling. However, these “noise”-induced curvatures were much smaller than those in the data (right column in the panel, *p* < 0.001). The plots are based on firing rates in the same window as in Fig. 2G, H (350-450 ms after stimulus onset). Colored lines are second-order polynomial fits to ten bootstrap samples of the observed data (left column) or ten simulated linearized population response manifolds (middle column). (**B**) Significant manifold curvatures were consistently observed in both monkeys performing the face task (monkey L, n = 62; monkey A, n = 70). (**C**) The curvature was also observed in individual sessions of the task when we could record enough units simultaneously to infer the manifold from single sessions (session 1, n = 8; session 2, n = 10). Conventions in B and C are the same as in Fig. 2G. (**D**) Average PSTHs of well-isolated single units (n = 75) and their curved manifolds showed the same patterns obtained from all units.

**Figure S6:**
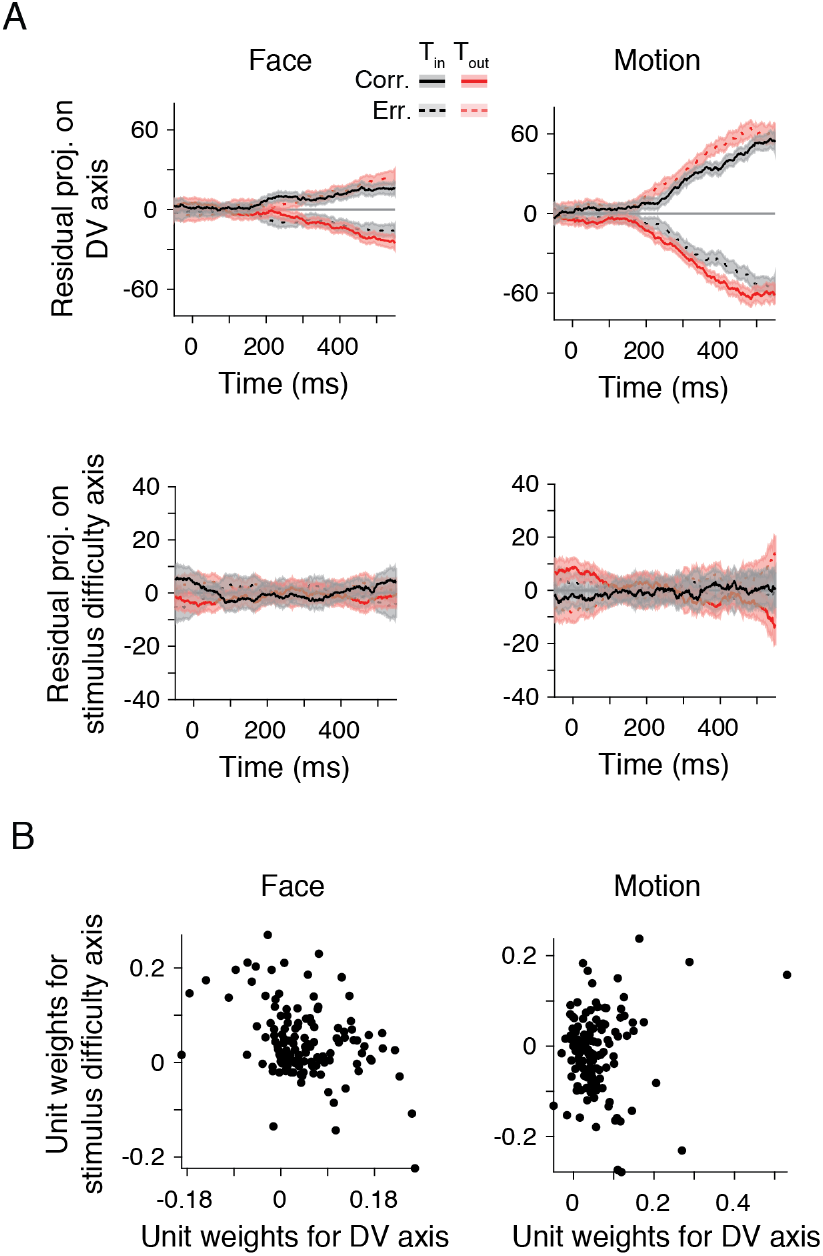
Properties of joint encoding of the DV and stimulus difficulty. (**A**) The monkey’s choices were correlated with projections of population responses along the DV axis but not along the difficulty axis. We calculated residual projections along each axis for each stimulus strength and choice by subtracting the mean projection for each condition. The plots show average residuals across stimulus strengths for each choice. Because we were interested in comparing correct and error choices here, we limited the analysis to the three lowest stimulus strength levels in each task, where error choices happened in the majority of sessions. In both tasks, the residual neural responses along the DV axis diverged depending on the monkey’s choices (T_in_ vs. T_out_, *p* < 0.001 in both tasks; 350-450 ms after stimulus onset, bootstrap test), whereas the residual responses along the stimulus difficulty axis were statistically indistinguishable (T_in_ vs. T_out_, face task, *p* = 0.15, motion task, *p* = 0.46). The absence of the correlation with behavioral performance is indicative that the activity along the stimulus difficulty axis does not reflect the monkeys’ increased task engagement or attention level. (**B**) The distribution of the weights of recorded units in the DV and stimulus difficulty axes of the face (n = 132) and motion tasks (n = 129). There was no apparent clustering of units preferentially encoding the DV or difficulty.

**Figure S7:**
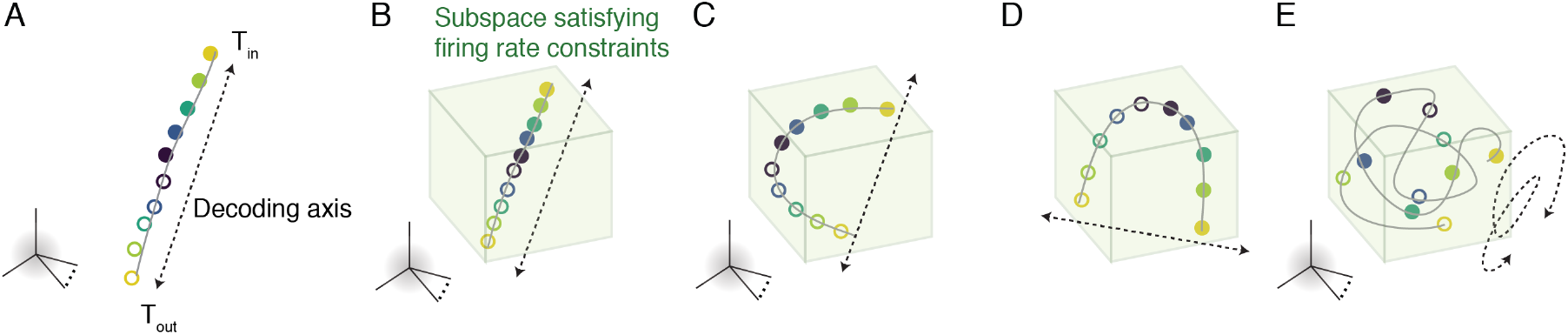
Curved manifolds emerge when precise coding of continuous variables must happen with a limited dynamic range of neural responses. Various constraints limit the dynamic range of neurons in a biophysically plausible neural circuit (Wohrer et al. 2013), including but not limited to the non-negativity of firing rates and metabolic costs associated with spiking (Keemink and Machens 2019; Lennie 2003). These constraints limit the neural responses to a bounded subspace within the population response state space. (**A**) Encoding a continuous variable (e.g., the DV) along a straight line in the state space can achieve arbitrary high precision in the absence of constraints on the dynamic range of neural responses. (**B**) Compacting a straight linear manifold within the bounded subspace has two important downsides. First, the diagonal lines that provide the maximum dynamic range go through the center of the bounded subspace, where the whole population has high levels of activity —an energetically expensive code. Second, compacting the straight manifold makes the response patterns associated with different DVs less distinguishable, reducing the encoding precision. (**C**) Bending the manifold within the bounded subspace will reduce energetic costs (Keemink and Machens 2019). Further, under a favorable correlation structure in the population responses (Moreno-Bote et al. 2014) and especially if decoding along a curved manifold is feasible, the curved manifold makes the population responses associated with neighboring DV values more distinguishable, increasing the encoding precision. (**D**) The presence of curvature is not changed by affine transformations of the bounding subspace, e.g., applying different scaling coefficients to the responses of individual units. However, the exact shape and layout in state space could be modified. (**E**) Although it is possible to further elongate the one-dimensional manifold by twisting and tangling it within the subspace, it may not be advantageous for two reasons. First, although such twisted manifolds allow larger separation of points along the manifold, they may bring distant points closer to each other in the state space, increasing readout errors due to noise. Second, a twisted manifold necessitates a more complex readout (decoder). Whereas a linear or low-order polynomial readout is easily implementable, more complex readouts may not be easy to implement or may lack the robustness offered by simpler decoders.

**Figure S8:**
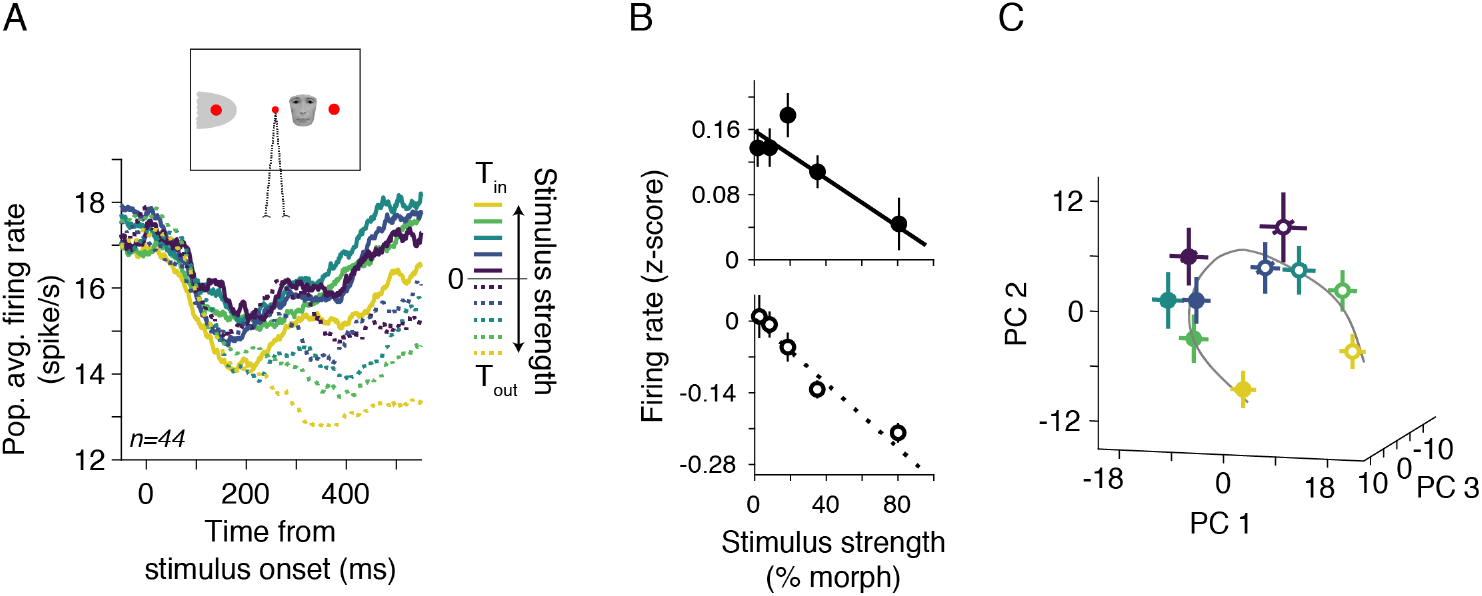
The results of a control experiment with face stimuli presented ipsilateral to the recording site were similar to the main task with contralateral stimuli. (**A**) Ipsilateral placement of face stimuli eliminated any possible overlap with the RF. Yet, the ordering of the PSTHs remained the same as that in the main task, confirming that the reversal of T_in_ PSTH order with stimulus strength was not caused by inadvertent overlaps between the RFs and the face stimuli. These results also rule out explanations based on attentional mechanisms or visual selectivity of LIP cells that depend on the overlap of the stimulus and the RF. Further support for our conclusions comes from independent analyses of stimuli with different sizes, which changed by an octave across trials. Both large and small stimuli in the main task caused consistent T_in_ PSTH orders and curved manifolds (not shown). Forty-four units were recorded in 17 sessions of the control experiment. Conventions are the same as in Fig. 2A. (**B**) The average firing rates in the control experiment decreased with stimulus strength for both T_in_ (Eq. 1, *β*_1_ = −0.15 ± 0.05, *p* = 0.010) and T_out_ choices (*β*_1_ = −0.31 ± 0.05, *p* = 4.6 × 10^−8^). (**C**) The population neural responses in the control experiment formed a curved manifold. The plot shows the neural activity at 400 ms after stimulus onset (100 ms window). The curvature of the manifold was significant (*p* < 0.001, based on tests of consistency of curvature direction, see Methods). Conventions are the same as in Fig. 2G.

**Figure S9:**
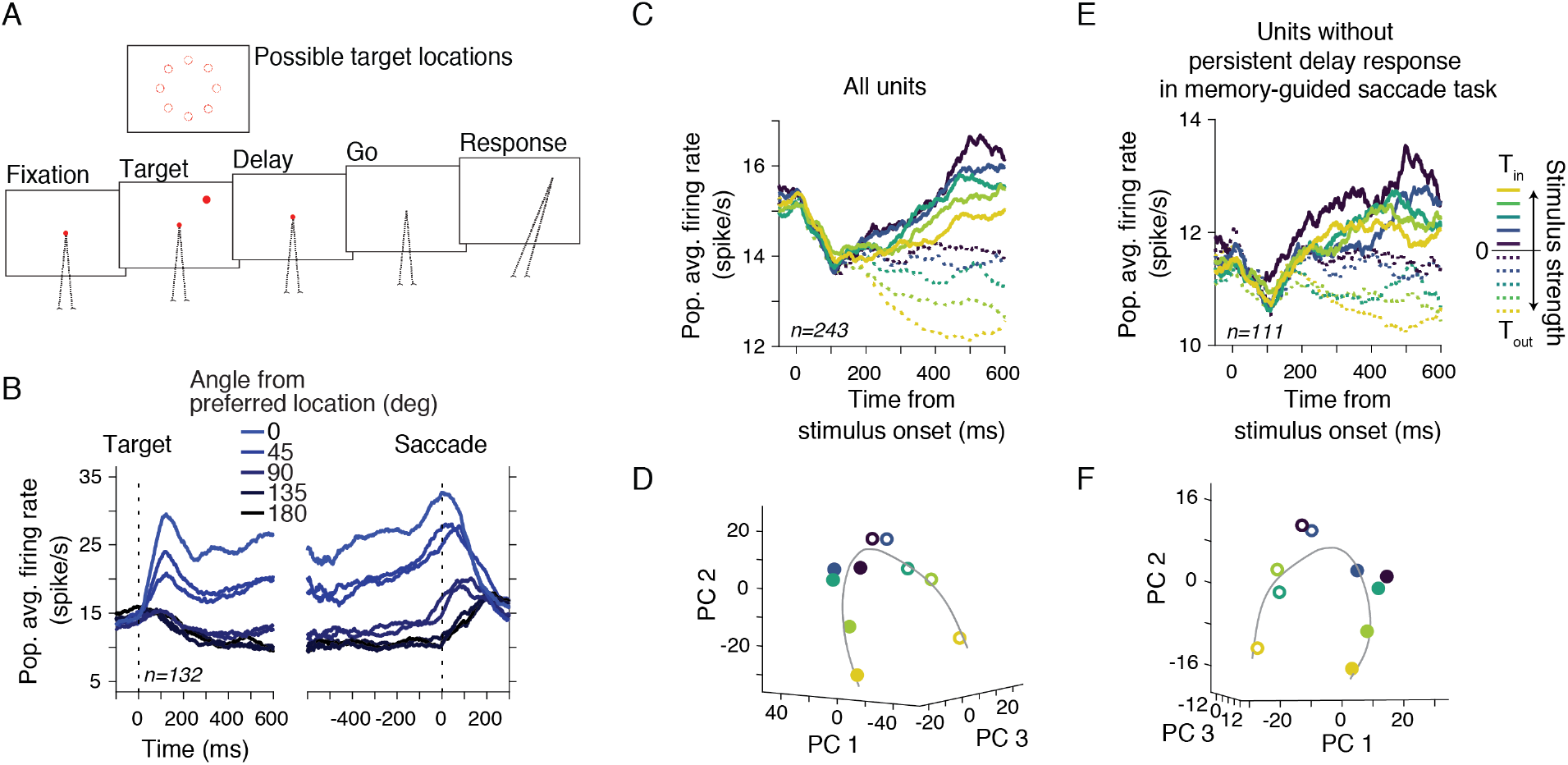
LIP neural responses during a memory-guided saccade task and their relevance to the DV encoding. (**A**) Task design. Prior to the face or motion tasks in each session, we performed a memory-guided saccade task to measure the RF location and size of the LIP neurons and confirm their sustained memory responses (Gnadt and Andersen 1988; Shadlen and Newsome 2001). In this task, after the monkey fixated on a central fixation point, a target flashed for 125 ms at one of eight possible locations on the screen. The monkey maintained fixation after the flash during a ~1000 ms delay period. At the end of the delay, the fixation point turned off, instructing the monkey to make a saccade to the remembered target location. The eccentricity of the target array was adjusted for each recording site based on manual mapping of RFs prior to the memory-guided saccade task. (**B**) Population average neural responses in the memory-guided saccade task. The figure includes all the units included in the main analyses of the face task. The preferred location was defined as the position in the target array that elicited the maximum firing rate during the delay period. Neurons showed consistent selectivity for their preferred location during visual, delay, and saccade periods, as typically observed in LIP (Gnadt and Andersen 1988). (**C-F**) Although our main analyses focused on neurons with significant persistent activity during the delay period of the memory-guided saccade task to remain consistent with past studies, the cells that did not elicit significant persistent activity in the saccade task showed similar population response profiles in our main tasks. Including these cells in our analyses for the face task (n=111) did not critically change the reverse ordering of PSTHs (C) or the curved manifold of the population responses (D). In fact, the population of the cells without persistent activity in the memory-guided saccade task replicated both findings on their own (E-F), albeit they had lower firing rates during the stimulus viewing period.

## Notes

### Competing Interest Statement

The authors have declared no competing interest.

